# Low cost and open source multi-fluorescence imaging system for teaching and research in biology and bioengineering

**DOI:** 10.1101/194324

**Authors:** Nuñez Isaac, Matute Tamara, Herrera Roberto, Keymer Juan, Marzullo Tim, Rudge Tim, Federici Fernán

## Abstract

The advent of easy-to-use open source microcontrollers, off-the-shelf electronics and customizable manufacturing technologies has facilitated the development of inexpensive scientific devices and laboratory equipment. In this study, we describe an imaging system that integrates low-cost and open-source hardware, software and genetic resources. The multi-fluorescence imaging system consists of readily available 470 nm LEDs, a Raspberry Pi camera and a set of filters made with low cost acrylics. This device allows imaging in scales ranging from single colonies to entire plates. We developed a set of genetic components (e.g. promoters, coding sequences, terminators) and vectors following the standard framework of Golden Gate, which allowed the fabrication of genetic constructs in a combinatorial, low cost and robust manner. In order to provide simultaneous imaging of multiple wavelength signals, we screened a series of long stokes shift fluorescent proteins that could be combined with cyan/green fluorescent proteins. We found CyOFP1, mBeRFP and sfGFP to be the most compatible set for 3-channel fluorescent imaging. We developed open source Python code to operate the hardware to run time-lapse experiments with automated control of illumination and camera and a Python module to analyze data and extract meaningful biological information. To demonstrate the potential application of this integral system, we tested its performance on a diverse range of imaging assays often used in disciplines such as microbial ecology, microbiology and synthetic biology. We also assessed its potential for STEM teaching in a high school environment, using it to teach biology, hardware design, optics, and programming. Together, these results demonstrate the successful integration of open source hardware, software, genetic resources and customizable manufacturing to obtain a powerful, low cost and robust system for STEM education, scientific research and bioengineering. All the resources developed here are available under open source licenses.

## Introduction

Fluorescence imaging has become an essential tool for research and education in biological sciences and bioengineering. However, instrumentation for fluorescence imaging is expensive and access to software is often restricted, which imposes a high barrier for democratizing its usage outside academic labs and wealthy research institutions. Open hardware and free/libre and open source software (FLOSS) provides the tools for fabricating these devices at a much lower cost and with better adaptability. The advent of easy-to-use microcontrollers, off-the-shelf electronics and customizable manufacturing technologies such as 3D printers and laser-cutting machines, has given rise to a diverse community of open hardware developers, users and tinkerers. This ethos has facilitated the development of inexpensive scientific devices and laboratory equipment [1–4]; such as PCR thermocyclers [5], photometric sensors [6], a quartz crystal microbalance [7] and neuroscience instrumentation [8–10], among many others [11]. The use of online platforms for project documentation and sharing [12–14], along with the implementation of collaborative practices borrowed from the FLOSS community, is enabling crowdsourced development of advance instrumentation (e.g [15]). These modes of development are lubricated by licenses that guarantee freedom to access, use, modify and distribute source code, design files and derived works.

A broad series of open source digital microscopes and imaging devices have been created (e.g. [16–20]). Some of these include advanced features such as light-sheet illumination [21, 22], two photon scanning [23], a motorized stage [24, 25], optogenetic stimulation [26], cell phone adaptability (e.g. [27–31]), wireless connectivity [32], and cost as little as US$1 [33]. Open source devices for in vivo fluorescence imaging of whole organisms and cell populations are of special interest for spatio-temporal studies [26, 34, 35]. Fluorescence imaging has traditionally relied on broad-spectrum illumination systems, such as mercury lamps, that require channel separation for multicolor imaging. This is usually achieved by switching filters, which involves moving mechanical parts that complicates designs and can cause unwanted vibrations in imaging mainframes. One alternative is to use multiband filters and switch between multiple Light Emitting Diodes (LEDs) whose output can be modulated in microseconds to excite different fluorophores consecutively [36].

Here, we combined low-cost color cameras, off-the-shelf electronics, inexpensive acrylic filters, open-source software, easy-to-use genetic resources, and recently developed long Stokes shift fluorescent proteins to create an integrated imaging system for multiple fluorescence signals from a single excitation wavelength. The device was designed to allow imaging at different scales, ranging from a single bacterial colony (~500-1000 μm) to entire petri dishes (~10 cm), scales not often covered by commercially available devices. We explored the applications and limitations of this system in a series of diverse experimental setups that required spatio-temporal tracking of multiple fluorescence signals. We demonstrate the value and robustness of our system for low cost STEM education, scientific research and bioengineering. All the resources developed here are available under open source licenses.

## Materials and Methods

### DNA construction

All vectors were constructed by Golden Gate using BsaI (NEB) and T4 ligase (NEB). The aceptor vectors for Golden Gate assembly were constructed by Gibson Assembly. Nine level 0 vectors and one acceptor vector compatible with the CIDAR MoClo Toolkit (Addgene) were constructed by Gibson Assembly. The CIDAR MoClo Parts Kit was a gift from Douglas Densmore (Addgene kit # 1000000059). The PCR fragments were amplified using Phusion High-Fidelity ADN Polymerase (NEB) and purified using Wizard SV Gel and PCR Clean-Up System (Promega). The purification of all plasmids was performed using Wizard Plus SV Miniprep DNA Purification System (Promega). All the assembled vectors were checked by colony PCR and later sequenced commercially (Macrogen Inc.). See Table 1 for full description of genetic parts. All the genbank files can be downloaded from the following Open Science Framework link: https://osf.io/dy6p2/.

**Table 1.** Genetic components used in this study

| Component | Type | Reference |
| --- | --- | --- |
| A_J23101_B | Level 0: promoter | This study |
| J23116_AB | Level 0: promoter | [46] |
| J23107_AB | Level 0: promoter | [46] |
| J23100_AB | Level 0: promoter | [46] |
| R0040_pTet_AB | Level 0: promoter | [46] |
| A_pLux76_B | Level 0: promoter | This study |
| A_pLas81_B | Level 0: promoter | This study |
| R0010_pLac_AB | Level 0: promoter | [46] |
| BCD2_BC | Level 0: RBS | [46] |
| BCD8_BC | Level 0: RBS | [46] |
| BCD12_BC | Level 0: RBS | [46] |
| B0032m_BC | Level 0: RBS | [46] |
| B0033m_BC | Level 0: RBS | [46] |
| B0034m_BC | Level 0: RBS | [46] |
| B_RiboJ54_C | Level 0: RBS | This study |
| C_mBeRFP_D | Level 0: CDS | This study |
| C_sfGFP_D | Level 0: CDS | This study |
| C_CyOFP_D | Level 0: CDS | This study |
| C_hmKeima8.5_D | Level 0: CDS | This study |
| C_mTurquoise2_D | Level 0: CDS | This study |
| C_mRuby2_D | Level 0: CDS | This study |
| C_mTFP1_D | Level 0: CDS | This study |
| C_luxI_D | Level 0: CDS | This study |
| C_lasR C0079_D | Level 0: CDS | This study |
| C_lasI_D | Level 0: CDS | This study |
| E0030_YFP_CD | Level 0: CDS | [46] |
| C0062_luxR_CD | Level 0: CDS | [46] |
| B0015_DE | Level 0: Terminator | [46] |
| D_ECK0818_E | Level 0: Terminator | This study |
| D_ECK9600_D | Level 0: Terminator | This study |
| pLac_RiboJ54_YFP_B0015 | level 1 vector used in Fig 2A | This study |
| pLac_RiboJ54_mRuby2_B0015 | level 1 vector used in Fig 2A | This study |
| pLac_RiboJ54_mTurquoise2_B0015 | level 1 vector used in Fig 2A | This study |
| pLac_RiboJ54_mBeRFP_B0015 | level 1 vector used in Fig 2A | This study |
| pLac_RiboJ54_hmKeima8.5_B0015 | level 1 vector used in Fig 2A | This study |
| pLac_RiboJ54_mTFP1_B0015 | level 1 vector used in Fig 2A | This study |
| pLac_RiboJ54_sfGFP_B0015 | level 1 vector used in Fig 2A | This study |
| pLac_RiboJ54_CyOFP_B0015 | level 1 vector used in Fig 2A | This study |
| J23116_BCD12_sfGFP_B0015 | level 1 vector used in S1 Fig | This study |
| J23107_BCD12_sfGFP_B0015 | level 1 vector used in S1 Fig | This study |
| J23100_BCD12_sfGFP_B0015 | level 1 vector used in S1 Fig & Fig 2E | This study |
| J23101_BCD12_sfGFP_B0015 | level 1 vector used in S1 Fig & Fig 2E | This study |
| J23101_RiboJ54_sfGFP_B0015 | level 1 vector used in Fig 2E | This study |
| J23101_RiboJ54_mBeRFP_B0015 | level 1 vector used in Fig 2E | This study |
| J23101_RiboJ54_CyOFP_B0015 | level 1 vector used in Fig 2E, 2B, 4, 6A | This study |
| J23101_BCD12_mBeRFP_B0015 | level 1 vector used in Fig 2E | This study |
| J23101_BCD12_CyOFP_B0015 | level 1 vector used in Fig 2E | This study |
| J23101_BCD8_sfGFP_B0015 | level 1 vector used in Fig 2E, 2B, 4, 6A | This study |
| J23101_BCD8_mBeRFP_B0015 | level 1 vector used in Fig 2E | This study |
| J23101_BCD8_CyOFP_B0015 | level 1 vector used in Fig 2E | This study |
| J23101_BCD2_sfGFP_B0015 | level 1 vector used in Fig 2E | This study |
| J23101_BCD2_mBeRFP_B0015 | level 1 vector used in Fig 2E | This study |
| J23101_BCD2_CyOFP_B0015 | level 1 vector used in Fig 2E | This study |
| J23101_B0034_sfGFP_B0015 | level 1 vector used in Fig 2E | This study |
| J23101_B0034_mBeRFP_B0015 | level 1 vector used in Fig 2E, 2B, 4, 6A | This study |
| J23101_B0034_CyOFP_B0015 | level 1 vector used in Fig 2E | This study |
| J23101_B0033_sfGFP_B0015 | level 1 vector used in Fig 2E | This study |
| J23101_B0033_mBeRFP_B0015 | level 1 vector used in Fig 2E | This study |
| J23101_B0033_CyOFP_B0015 | level 1 vector used in Fig 2E | This study |
| J23101_B0032_sfGFP_B0015 | level 1 vector used in Fig 2E | This study |
| J23101_B0032_mBeRFP_B0015 | level 1 vector used in Fig 2E | This study |
| J23101_B0032_CyOFP_B0015 | level 1 vector used in Fig 2E | This study |
| 12X | level 1 acceptor vector | This study |
| 1LU2 | vector used in Fig 3 | This study |
| plac34lasI | vector used in Fig 3 | This study |
| plac34luxI | vector used in Fig 3 | This study |
| plac54lasI | vector used in Fig 3 | This study |
| plac54luxI | vector used in Fig 3 | This study |
| plas8O | vector used in Fig 3 | This study |
| plas33O | vector used in Fig 3 | This study |
| plux34G | vector used in Fig 3 | This study |
| plux54G | vector used in Fig 3 | This study |
| ptet32LasR | vector used in Fig 3 | This study |
| Std34BeRFP | vector used in Fig 3 | This study |

### Growth conditions

All tests were carried out on *E.coli* Top10 (Invitrogen) grown at 37°C in LB medium. Media were supplemented with 1.5% m/v agar for assays on solid media. For inducer applications, we prepared a stock solution of 0.1 M IPTG, sterilized by 0.22 mm filter. Antibiotics were used as solutions of kanamycin (50 ug/ml) and carbenicillin (100 μg/ml).

Different fluorescent proteins were tested by streaking transformed cells onto petri dishes from glycerol stocks and growing them for 24 hrs at 37°C. Expression was induced with 30 μl of 0.1M IPTG.

### Transformations and co-transformations

The chemocompetent cells used for all the transformations were made by the CCMB80 method in *E.coli* Top10 (Invitrogen) (OpenWetWare: "TOP10 Chemically competent cells", 2013). Cells stored at −80 ° C were thawed on ice for 20 minutes. For single transformations, a 1 μl of plasmid stock (concentration ~ 1ng/μl) was added to 50 μl of chemocompetent Top10 *E. coli.* In the case of co-transformations, 100 ng total of each plasmid were mixed with deionized water to obtain a final volume of 6 μl, which were then added to 50 μl of chemocompetent Top10 *E. coli*. The cells were incubated for 25 minutes and heat-shocked at 42°C for 1 minute, after which they were incubated on ice for an additional 2 minutes.

A 250 μl volume of LB medium was added to each transformation and incubated with shaking (250 rpm) at 37°C. The incubation time for the single transformation was 1 hour, after which a volume of 25 μl of transformed cells was plated in solid LB medium supplemented with antibiotics. In the case of co-transformations, the incubation time was 1.5 hours, after which 35 μl of transformed cells were plated on LB-agar supplemented with antibiotics. Plates were incubated at 37°C overnight (~ 12-16 hours) for colony formation.

### Colony-sectoring assays

Cultures of the three TOP10 cells containing the different vectors were grown overnight and mixed proportionally. The culture mixture was diluted 1:100 or 1:10,000 and a 0.5 μl drop was placed in the center of a petri dish. Plates were then incubated at 37°C for 12 hours and imaged every 12 hours for 7 days at room temperature.

We have created a protocol online describing this experimental procedure in more details (dx.doi.org/10.17504/protocols.io.jsecnbe).

### Grown-on-membrane assays

An ISO-GRID membrane filter (Neogen corporation, 1600 Grid 0.45μm, 36 Grid; product #6802) was placed on top of LB agar plates, with the hydrophobic grid facing up. Bacteria co-transformed with the sender/receiver plasmids were grown at 37°C in LB liquid media to OD600 0.03 (~5 hours) and 0.5 μl of cell culture was pipetted within each well following arrangements that depend on the assays. We have created a protocol online describing this experimental procedure in more details (dx.doi.org/10.17504/protocols.io.jsfcnbn).

### Mix colonies time lapses

Time lapses and experiments shown in Fig 4 and 5 were performed by mixing liquid cultures of the different strains transformed with mBeRFP, CyOFP1 and sfGFP (eg, 100 μl of each one), performing three consecutive 1:100 volume dilutions of the culture (pure or mixture) and plating this mixed culture on a LB agar plate. We have created a protocol online describing this experimental procedure in more details (dx.doi.org/10.17504/protocols.io.jsucnew).

### Camera settings

For Fig 2 and Fig 3 the camera was operated using the command raspistill as follows:

**Fig 1.**
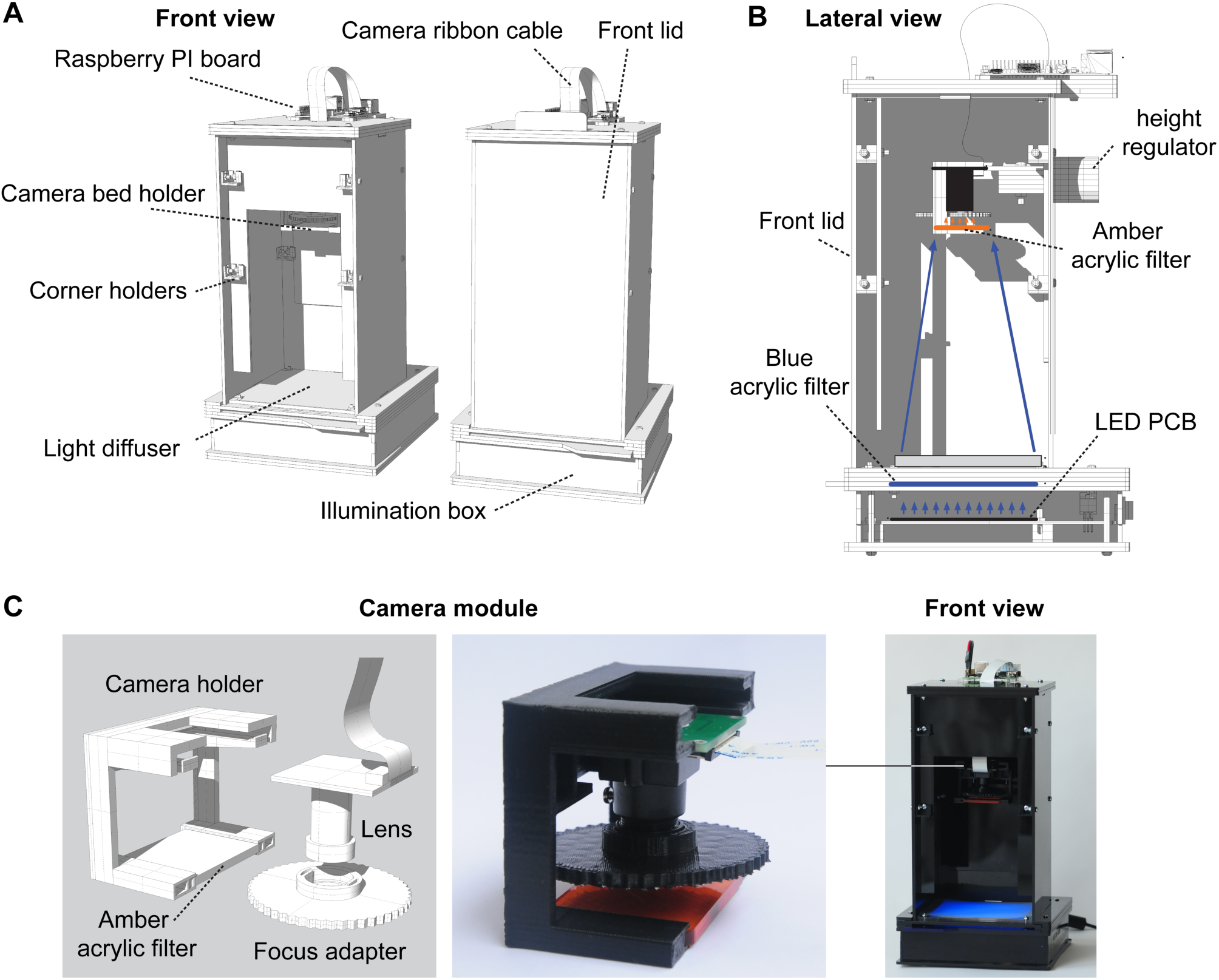
Device general architecture. (A) Schematic rendering of the device with and without the front lid. (B) lateral view of a longitudinal cross-section of the device showing height control, illumination box and filters. (C) Camera and amber filter holder.

**Fig 2.**
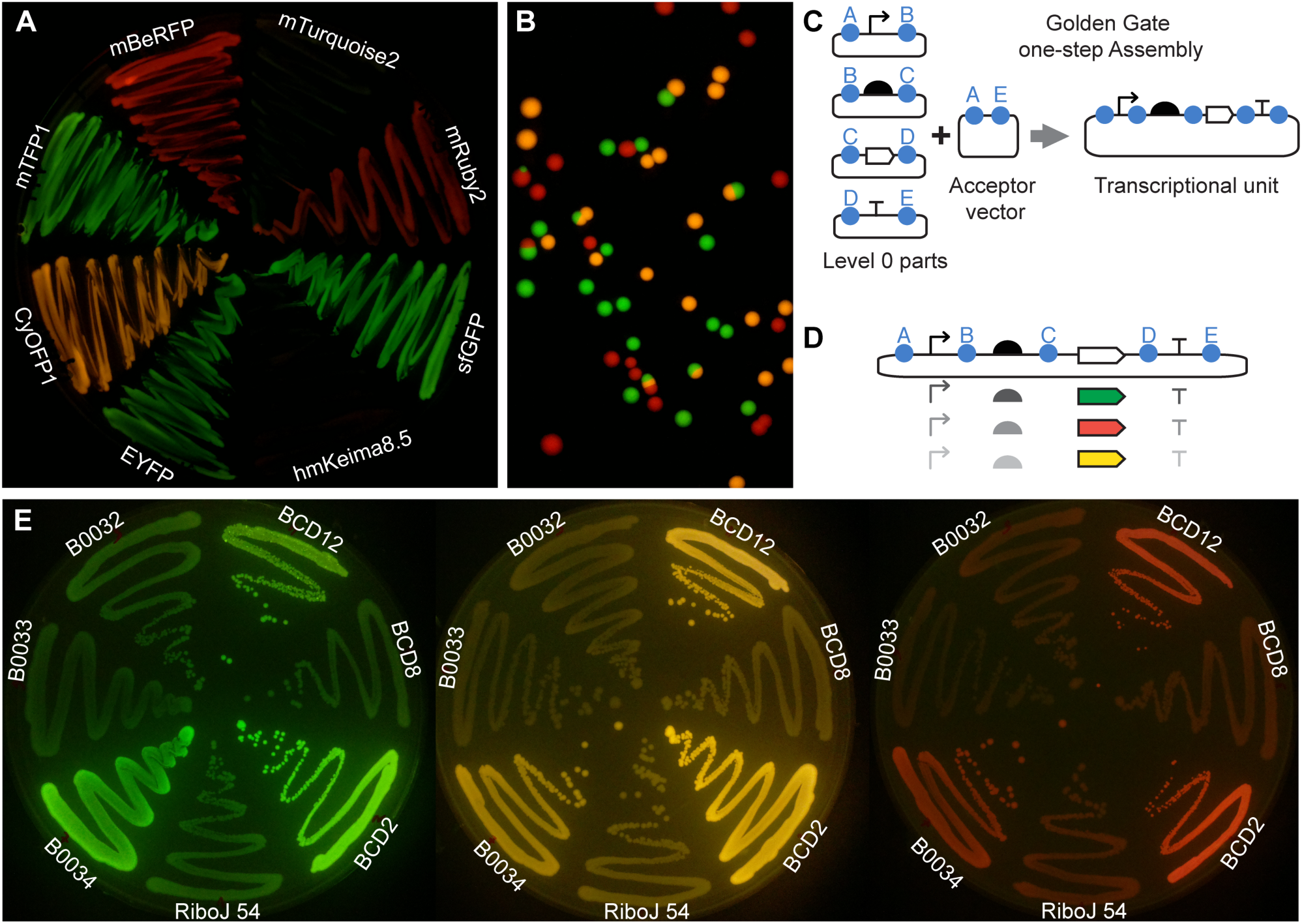
Simultaneous imaging of different fluorescent proteins with single excitation at 470 nm. (A) CyOFP1, hmKeima8.5, mBeRFP, mRuby2, mTFP1, sfGFP, mTurquoise2, and EYFP, with R0010 promoter and RiboJ 54 insulator-RBS treated with (right) and without (left) IPTG. (B) Representative image showing growing E. coli colonies expressing sfGFP, CyOFP1 and mBeRFP. (C) Schematic representation of Golden Gate one-step assembly for the construction of genetic systems from level 0 parts. (D) Schematic representation of combinations of promoter, RBS, CDS and terminator that can be assembled with the standardized Golden Gate method. (E) Image of agar plates with streaks of *E. coli* expressing sfGFP, CyOFP1 and mBeRFP under J23101 promoter and different ribosome binding sites (RBS): BCD12; BCD8; BCD2; RiboJ54; B0034; B0033; and B0032. B0015 terminator was used for all the constructs. No post-editing has been applied to these images; only cropping and alignment.

**Fig 3.**
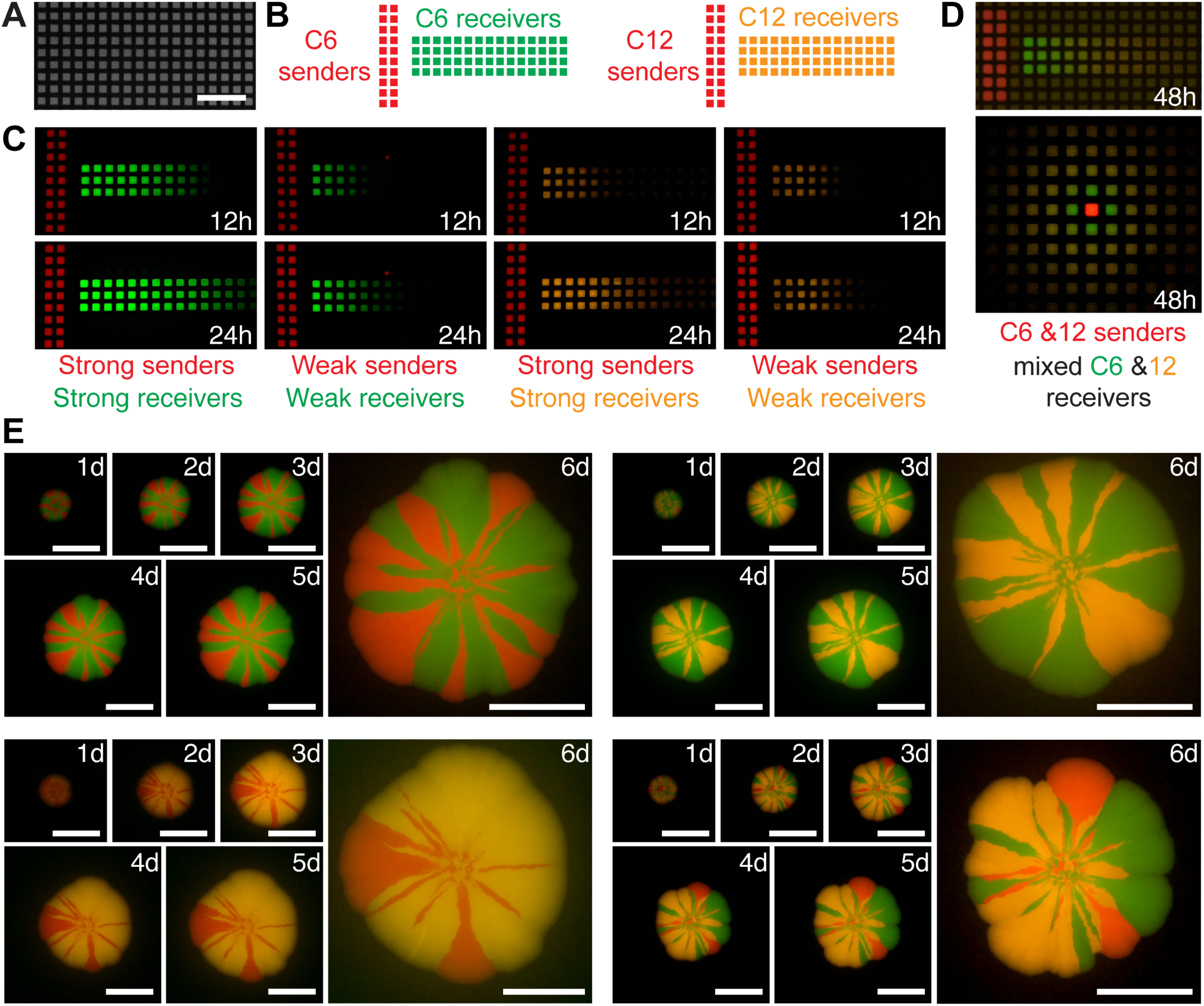
System testing under different experimental setups. (A) Image of ISO-GRID permeable membrane used in experiments shown in B to D. Scale bar, 1000 μm. (B) Schematic representation of diffusion assay for *E. coli* cells producing (“sender cells”) and responding to (“receiver cells”) C6 and C12 homoserine lactones plated on permeable membranes printed with hydrophobics grid lines. (C) Images of a time-lapse experiment of cell responses to C6 and C12 homoserine lactones. Images were taken at 12 and 28 hours after plating. Cells were grown at 37°C outside the device. Vector co-transformation for: strong C12 receiver, pTet_32 LasR + pLas8O; weak C12 receiver, pTet_32 LasR + pLas33O; strong C6 receiver, 1LU2 + pLux34G; weak C6 receiver, 1LU2 + pLux54G; strong C6 sender, Std34BeRFP + pLac34LuxI; weak C6 sender, Std34BeRFP + pLac54LuxI; strong C12 sender, Std34BeRFP + pLac34LasI; weak C12 sender, Std34BeRFP + pLac54LasI. See S1 table for sequence information. Scale bar, 1000 μm. (D) Images at 48h after plating a mix of C6 and C12 sender cells (expressing mBeRFP, in red) and a mix of C6 and C12 receiver cells expressing sfGFP and CyOFP, respectively. ROH: pTet_32 LasR + pLas8O. (E) Colony-sectoring assay. Strains labelled with CyOFP1, mBeRFP and sfGFP fluorescent proteins seeded on agar LB plates in combinations of two (CyOFP1-mBeRFP, CyOFP1sfGFP and sfGFP-mBeRFP) or three (CyOFP1, mBeRFP and sfGFP) and tracked every 24h for 6 days. No post-editing has been applied to these images; only cropping and alignment. Scale bar, 500 μm.

Fig 2A: raspistill -t 4000 -ss 90000 -ISO 300 -awbg 1,1 -co 50 -o

Fig 2B raspistill -t 5000 -ss 3000 -o

Fig 3A to D: raspistill -t 4000 -ss 100000 -ISO 300 -awbg 1,1 -co 60 -o

Fig 3E: raspistill -t 4000 -ss 70000 -ISO 300 -awbg 1,1 -co 40 -o and raspistill -t 4000 -ss 50000 -ISO 300 -awbg 1,1 -co 30 -o

Time lapses and experiments shown in Fig 4 and 5 were performed with python code controlling the camera as described in text.

**Fig 4.**
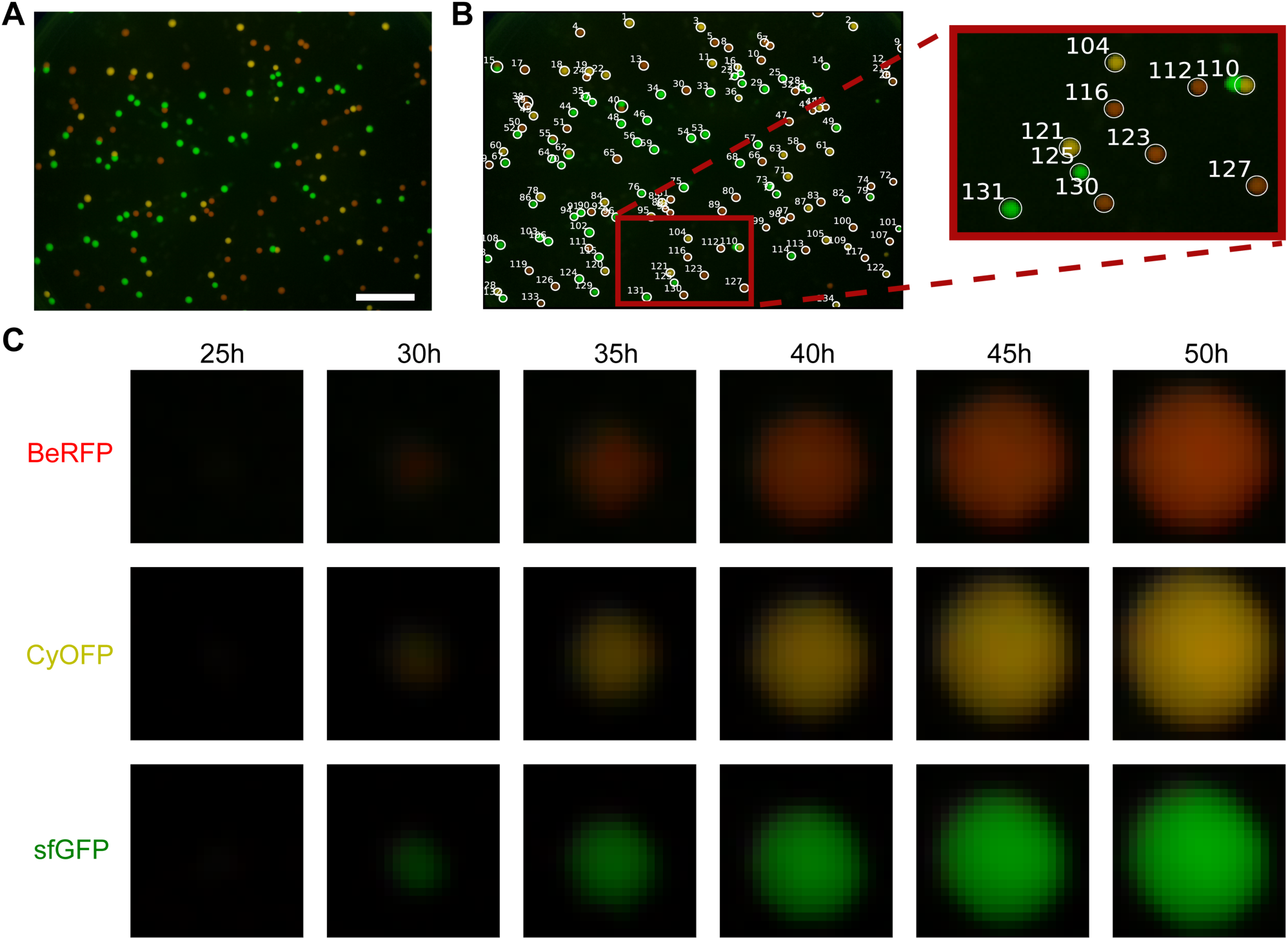
Colony identification and segmentation from timelapse images. (A) Endpoint timelapse image (see S4 movie) of a plate with growing colonies of three different strains, each one expressing a different fluorescent protein. Scale bar = 1cm. B) Colony identification (label them with an ID), by getting position in the image and final size. C) Segmentation of regions of interest (ROIs) defined by the standard deviation of the identified blobs. One example of each strain is shown.

**Fig 5.**
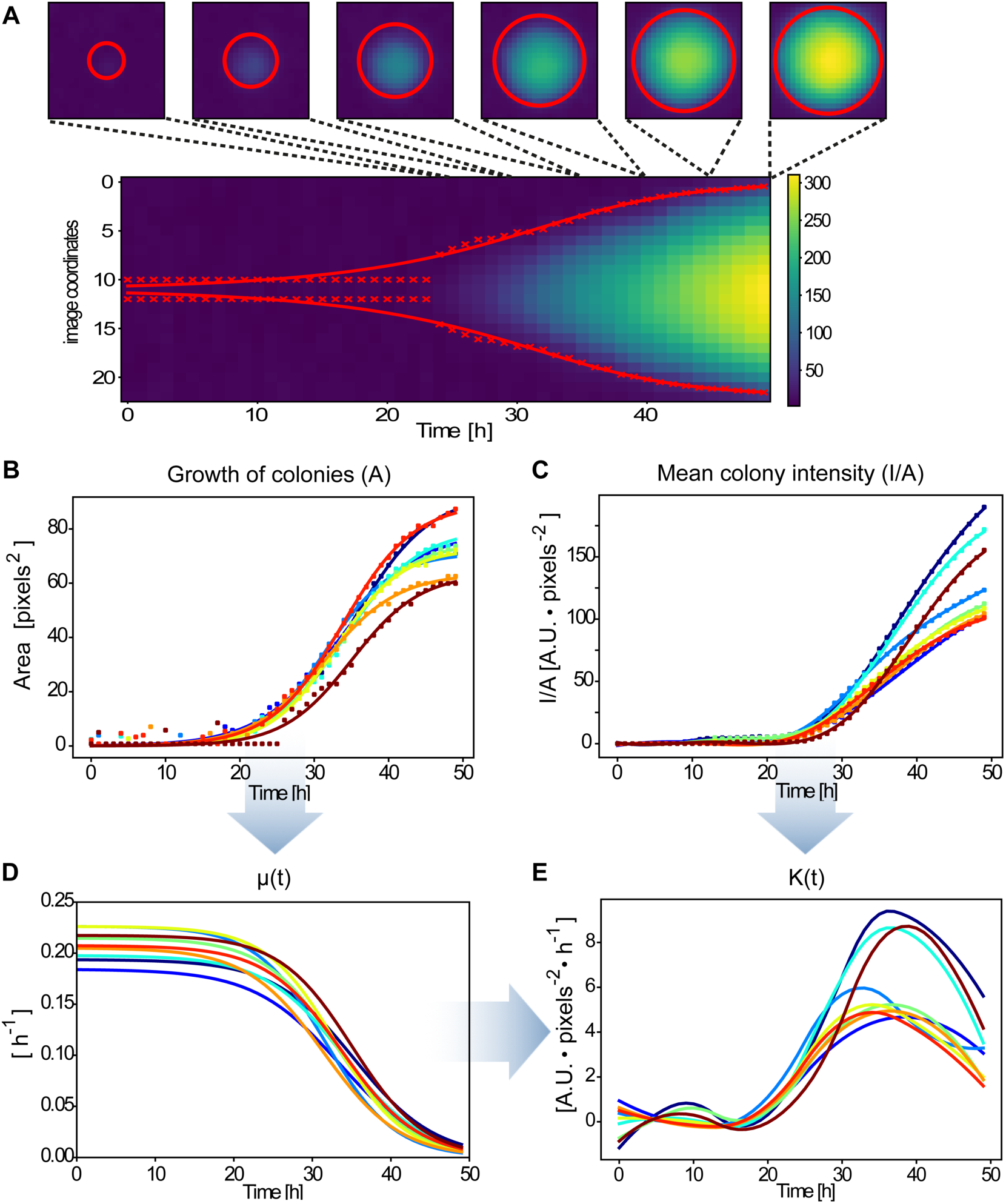
Analysis and parameter estimation from identified colonies. (A) kymograph of the pixel signal in the central slice of an example colony. The x marks show the radius estimated for each time and the red line is the radius value obtained from a fitted area function (also red circles). (B) Colony area curve fitting for some example colonies. Dots are the estimated area from the standard deviation of the blobs and the continuous lines are the model fitting of this values (C) Mean fluorescence intensity signal over time for the example colonies. Lines were obtained by spline smoothing. (D) Colony growth rate estimation from the data for each selected colony. (E) Protein expression rate parameter estimation using the previous data.

## Results

### Hardware Design

To allow for full customization, the mainframe was developed as enclosures made of editable laser-cut 3mm acrylic pieces (S1 Appendix). These can be easily stacked and shipped since they are mostly flat. The imaging device was assembled by holding together the acrylic plates with small 3d printed corner holders, designed to be used with standard nuts and bolts that can be replaced depending on local supply. All 3D printed parts were designed considering the FDM process: using mostly horizontal and vertical faces, low inclination angles and no “ceilings”, therefore no support structures were needed. The object architecture is fully modular (S1 Appendix).

The top module supports the Raspberry Pi for camera control (Fig 1A) (S2 Appendix). The top enclosure also provides support to the camera bed, height regulator and camera holder (S3 Appendix). The camera holder also supports an acrylic emission filter in front of the lens and a focus adapter wheel for manual adjustable focus lens (Fig 1B, 1C). Recorded images are transmitted to the Raspberry Pi with a standard 15 pin (300 mm) ribbon cable. Height adjustment works with a standard bolt and nut mounted with a knob, which slides up and down tightens for fixation on the back plate of the module. The maximum height of the imaging system allows the camera module to get images as wide as the illuminated area at the base of the top module, designed to fit a 90mm standard petri dish. The adapter wheel at the lens permits correcting for focus when height is changed. Together, the top enclosure maximizes the use of space while providing light-blockage for complete darkness during imaging (Fig 1C). The 3d printed adaptor that encloses the camera module is detachable, allowing for standalone use, transforming the camera in a powerful microscope and visualizer by its own, not described in this project.

The bottom module, inspired by open source blue transilluminators used for DNA electrophoresis gels [37], holds the illumination system composed of a matrix of 100 blue (470 nm) 5 mm LEDs. We designed a printed circuit board (“PCB”) to operate these LEDs, which also supports components for analog and digital on/off control (S4 Appendix). For digital control, a TIP122 transistor was used, considering 3.3 volts for the Raspberry GPIO control. A regulable 12-24 volts power supply was chosen so the PCB required less components (using 5 LEDs in parallel for a resistor) and there would not be a need for additional security requirements (i.e. a fuse). The excitation filter is housed in between the top plates of this enclosure and is fastened between the top adjusting screws and the main mounting screws on the base (Fig 1B). This is a blue 3 mm acrylic that slides in from the front, allowing the filter to be exchangeable. The distance between this blue filter and the LEDs was optimized to obtain an even distribution of light on the diffuser that sits at top of the bottom module. Care should be taken to ensure an even distribution of light if redesign of the bottom module is needed (e.g. varying the height between LEDs and diffuser/blue acrylic). The top module mounts on the bottom module by sliding in from the front and secures with the top adjusting screws. For detailed assembly instructions and full description of the pieces please see http://docubricks.com/viewer.jsp?id=701517893260717056.

### Genetic resources for simultaneous imaging of multiple fluorescent signals under single excitation/emission setup

In order to obtain multiple fluorescent signals simultaneously from a single excitation wavelength and a fixed filter, we evaluated a series of fluorescent proteins potentially excitable at 470 nm and detectable with the amber filter (Fig 2A). We tested CyOFP1 [38], hmKeima8.5 [39], mBeRFP [40], mRuby2 [41], mTFP1 [42], sfGFP [43], mTurquoise2 [44], EYFP [45]. We took images at different camera settings and evaluated their performance. For full transparency, none of the images shown in this work was edited. We selected sfGFP (ex 485/em 510) and the long stokes shift fluorescent proteins CyOFP1 (ex 497/em 589) and mBeRFP (ex 446/em 611) since they showed distinguishable signals in colonies of *Escherichia coli* grown in the same plate (Fig 2B). mTurquoise2 and hmKeima8.5 did not show detectable signals (Fig 2A).

All these fluorescent proteins, as well as all regulatory sequences (e.g. promoters, RBSes, terminators), developed in this work were created as modular parts standardized for fast, combinatorial and efficient fabrication of genetic constructs. In particular, we domesticated these components as “level 0” components following the CIDAR MoClo standards for Golden Gate assembly DNA assembly [45] (Fig 2C). This design also made our resources compatible with hundreds of parts distributed in other registries and labs (e.g. [46]). This method allowed us to combine the chosen fluorescent proteins with different regulatory elements (Fig 2D). We were able to test different ribosome binding sites (Fig 2D) and promoters (S1 Fig).

In order to demonstrate the range of possible applications of our system, we tested different experimental setups using the Raspberry Pi camera software (RPI foundation). First, we tested the detection of spatio-temporal fluorescent signals as a consequence of artificial cell–cell communication based on 3-oxo-C6- and 3-oxo-C12 homoserine lactones. Bacterial cells that produce (i.e. senders) and respond to (i.e. receivers) homoserine lactones were plated within wells outlined by hydrophobic ink on permeable membranes (ISO-GRID membrane filters) (Fig 3A-D). These grid membranes were placed on top of agar plates containing growth media (Fig 3A). This setup allows for cell containment in each well while permits the diffusion of the homoserine lactones through agar to neighboring wells (Fig 3B). Then, cellular responses to diffusive signals were tracked over time [47].

In order to provide resources that permit full flexibility in the construction of cells that produce and/or respond to these signals, we created level 0 parts for: i) LuxI and LasI enzymes, in charge of the synthesis of C6 and C12 HSLs, respectively. ii) LuxR and LasR genes, in charge of binding and responding to C6 and C12 HSLs, respectively. and iii) pLux and pLas promoters, from which transcription is induced by LuxR/LasR in response to C6-HSL or C12-HSL respectively. With these tools, we created strong and weak senders and receivers of both signals and registered their behaviour in time (Fig 3C). The imaging system showed good contrast and permitted to track CyOFP1, mBeRFP and sfGFP fluorescent signals simultaneously. Moreover, this system allowed us to track and distinguish the responses of mixed populations of C6 and C12 senders/receivers (Fig 3D).

Next, we evaluated an experimental design used in microbial ecology studies, colony-sectoring assay, in which a mix of bacterial strains is inoculated as a single drop on a petri dish and the dynamics of their growth registered in time lapse [48, 49] (Fig 3E). Strains labelled with CyOFP1, mBeRFP and sfGFP fluorescent proteins were plated in pairs and triplets. These plates were grown in a 37°C incubator and imaged every 24h over 6 days.

Since the device is modular and customizable, we were able to use a shorter top module for the last image at 6 days that permitted a closer position to the plate (see hardware description). The system permitted good detection and discrimination of the three signals. All images were taken with the same settings (described in materials and methods). Under these standard conditions, the CyOFP1-mBeRFP combination showed some background noise in the last image. None of these images has been edited, with the aim of showing raw data produced by our system. These data demonstrate the utility of our system for experimental setups commonly used microbiology, synthetic biology and microbial ecology studies.

### Open-source software for hardware control and data analysis

We developed and made available via github (www.github.com/synbiouc/fluopi), simple python code to run time-lapse experiments and store resulting image sequences (S1-S3 Movie). We also created an extensive Python module (fluopi) with functions for analyzing and visualizing these image sequences. We make extensive use of existing open-source projects Numpy, Scipy, Matplotlib, and Scikit-image, as well as the Raspberry Pi foundations python modules.

Separately we provide example Jupyter (IPython) notebooks containing code for analysis of time-lapse image series that contain detailed explanation of the analysis process. A simple tutorial notebook introduces the concepts used in loading images and processing them to extract biologically meaningful information. We also provide a simple tutorial notebook describing how to control the imaging hardware from Python. *Jupyter* provides an intuitive and interactive way to run and develop Python code while immediately viewing the results.

As such, these notebooks are ideal for teaching classes or as an introductory tutorial for those students new to programming.

Below we show how this open source software is used to find the location of colonies, estimate their size, growth-rate, and fluorescence and from these data extract the rate of gene expression.

### Images data management and colony slicing

Timelapse image sequences of bacterial colonies expressing fluorescent proteins contain valuable data about growth, gene expression and possible inter-colony interactions. The first step in extracting this information is to process images to generate separate signals for each colony’s size and fluorescence.

The images are composed of three channels - red (R), green (G), blue (B) -, each of which can contain different information about the cells and is managed separately. We created a Jupyter notebook to analyze timelapse images of growing colonies constitutively expressing different fluorescent proteins (Fig. 4A, S4 Movie) as an example of the kind of data it is possible to obtain with FluoPi and the fluopi Python module. We also provide a simplified tutorial demonstrating the ideas and maths behind the Python module (www.github.com/synbiouc/fluopi).

The first steps in the analysis involve data management, essential for teaching programming: reading the image files and organizing their data in arrays. Next image processing is introduced to remove the background signal, sum all the channel values over time for each pixel, and smooth the data with a gaussian filter. After this pre-processing a gaussian shape (“blob”) detector from *skimage* is used to find the position of each colony in the plate and assign labels to them (Fig. 4B). With the standard deviation of these blobs it is possible to define square regions of interest (SROIs) where colonies are situated (Fig. 4C) and allows management of each of them separately.

### Growth and fluorescent expression rate of colonies

From the data extracted from images as described above, it is possible to estimate the growth rate of each colony and the average constitutive gene expression rate of a fluorescent reporter, considering multiple colonies on a single plate. To get growth information - size (area) and growth rate (relative change in area) -, it is possible to approximate the colony size by the region expressing fluorescent signal. This size can be obtained by applying the gaussian blob detector to each SROI in all frames (i.e. time points), obtaining the colony radius/area over time (Fig. 5 A). The result is noisy, mainly at the beginning, and so we fit the sigmoidal area model in eq.(1) to the area values computed with the above radii (Fig. 5B).

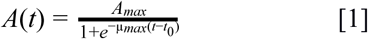

Then, as growth rate is defined by eq. (2), we are able to directly estimate the growth rate of the colonies with eq. (3) as shown in Fig. 5D

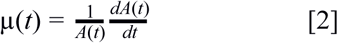

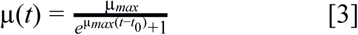

On the other hand, we show (see S5 Appendix) that the fluorescent expression rate, proportional to protein expression rate, is given by:

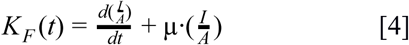

Then, we need to compute the mean intensity signal per unit of area (I/A). To achieve this, we get the colony fluorescent intensity in each frame by summing the three channel values for each pixel inside its computed area. Then dividing by the total number of pixels within this area it is possible to get the mean fluorescent intensity in each time step (Fig 5C). These values were smoothed by applying a smoothing spline function (scipy.interpolate.UnivariateSpline), which allows to compute analytically its derivatives with respect to time. Finally, with all these values, we are able to characterize the fluorescent protein dynamics by estimating the fluorescent reporter expression rate (Fig 5E).

### Distinguishing multiple fluorescent proteins from R,G,B color channels

An interesting aspect of being able to detect diverse fluorescent proteins is the possibility to label and identify different strains expressing them. To do this we first extract colony fluorescence signals as described above. With this data it is possible to get the characteristics of each fluorescent protein and classify the colonies according to strain.

These signals are in three channels, nominally labelled “red”, “green” and “blue” and depend on the filters and sensitivity of the camera. Each fluorescent protein has a characteristic spectrum which overlaps with these three channel spectral sensitivities. If we assume a linear relation between fluorescent protein expression and intensity of signal, it is possible to find a characteristic linear relation between the intensities in the (R,G,B) channels of each fluorescent protein (see “Colony classification maths” on S6 Appendix). That is, the relative intensity of each fluorescent protein in the (R,G,B) channels is distinct.

To test this, we performed independent timelapse experiments of strains expressing sfGFP, CyOFP and BeRFP (Fig 6A, S5-S7 Movie), from which we computed a characteristic linear relation for the emission of each protein in the red and green camera channels (Fig 6B). To test the capability of classification of this approach, we performed another timelapse with a mix of the three strains and classified each colony accord the closest characteristic linear relation between red and green camera channels (Fig 6C) (S8 Movie). This classification could be a useful tool to compute and compare attributes between groups of strains especially for microbial ecology studies.

**Fig 6.**
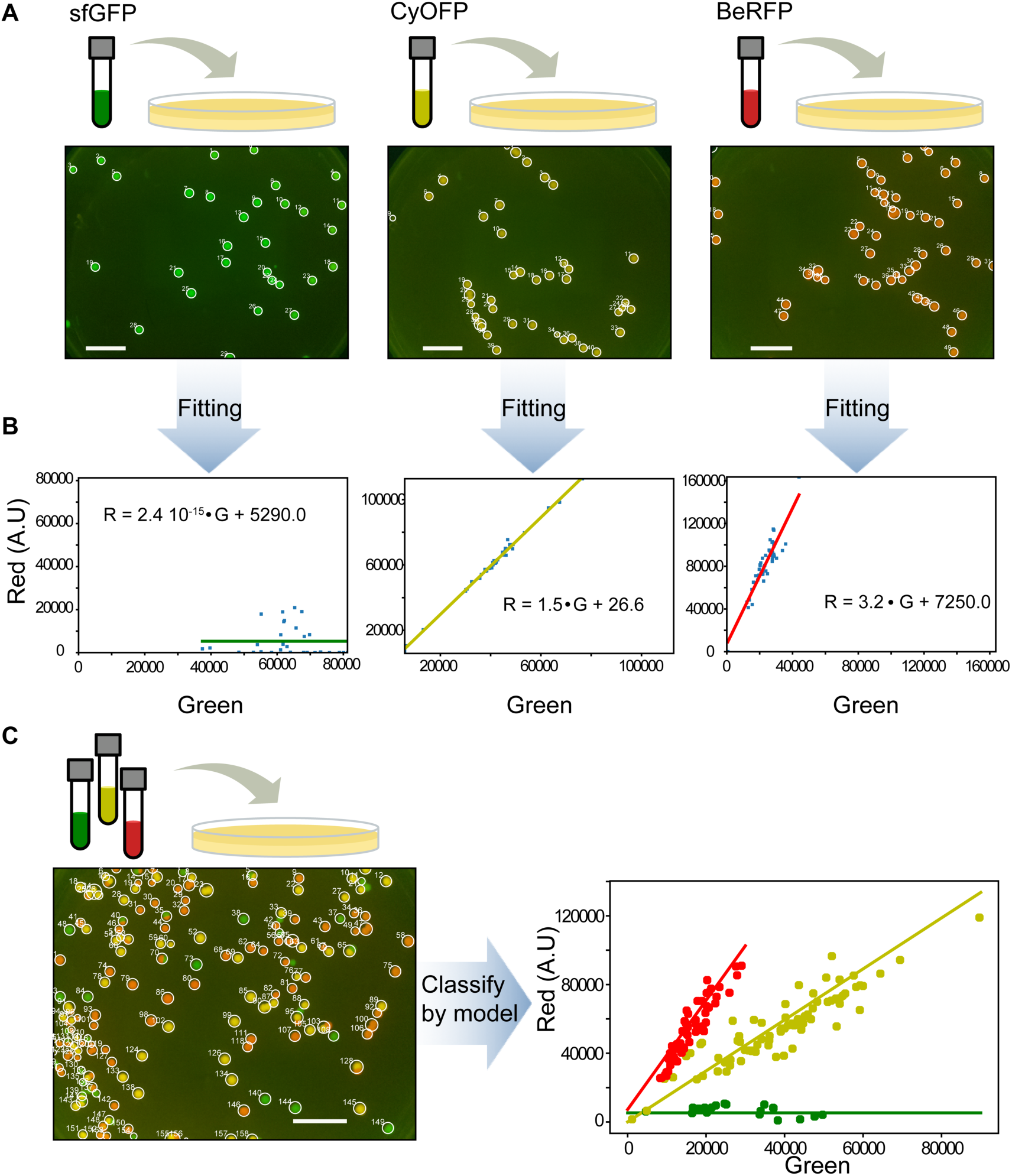
Distinguishing different strains by (R,G) color profile. (A) Timelapse images were taken separately of three strains, each expressing a different fluorescent protein (sfGFP, BeRFP and CyOFP). Figure shows final image (see S5-S7 Movie) with the detected colonies labeled. Scale bar = 1cm. (B) Characteristic linear relationship between red and green channel for each strain/fluorescent protein obtained from each timelapse in A. Each dot is the (R,G) total signal pair of a colony and the straight line is the linear regression. (C) When grown together on the same plate, the characteristic profile obtained could be used to classify the colonies according to strain/fluorescent protein (see S8 Movie). Scale bar =1 cm.

## Discussion

Following the impact of FOSS in software development, open hardware aims to democratize and accelerate hardware development in a similar manner. The scientific community has started to embrace this approach; with hundreds of labs around the globe designing and making their own equipment (e.g. [50]). Often these designs replace commercial scientific instrumentation, and self-manufacturing can reduce costs by 90-99% [4, 51], resulting in hundreds to thousands percent return on investment in science funding [52]. The total cost of our device is approximately US$250, including all components, materials, 3D printing and CNC cutting service fees. Commercial equivalents can cost as much as US$10000 with less functionality. Reducing costs and using accessible components lowers the access barrier to scientific experiments and democratizes science. In this way, open hardware engages talented developers and experimenters from non-traditional environments such as those in high schools, DIY movements, maker spaces, biohacking communities and art collectives (e.g. [14] [53–57]).

The open source nature of our project aims to contribute to and benefit from these non-traditional scientific communities, especially from the DIY microscopy community (e.g. [58]). To facilitate this process, we have provided full documentation of assembly steps (http://docubricks.com/viewer.jsp?id=701517893260717056) and operational code (github.com/synbiouc/fluopi). Good documentation is critical for crowdsourced design improvements. It also permits adaptations to local needs and restrictions, for example due to availability of parts, and facilitates the incorporation of new parts adapted from other open source projects. For instance the use of an adjustable focus RPI camera, instead of a fixed lens mounted on the Raspberry PI V2 camera, was suggested to us by another group working on similar developments (FlyPi microscope; [26]).

The imaging device is fully modular (S1 Appendix), permitting the re-engineering of parts separately. For instance, a shorter top enclosure that enabled imaging at much closer distances to the sample (while maintaining the full petri dish imaging capabilities) can be developed (github.com/synbiouc/fluopi). For instance, this shorter top enclosure was used for the 6-day colony images shown in Fig 3E. This was developed mainly due to restrictions on the length of the CSI camera cable provided by the manufacturer (and incompatibilities with longer cables found in the market). An alternative to this would be mounting the adjustable focus M12 lens on a 3D-printed holder [59] designed for the Raspberry PI V2 camera sensor, which is compatible with longer cables. This would also allow increasing the image resolution from 5Mp to 8Mp. Our modular design approach will also help in accommodating further improvements such as multi-wavelength LED fast multicolor illumination switching [36], temperature control, or automated focus control [26].

Our project integrates software, hardware and wetware to achieve single-excitation multiple-emission fluorescence microscopy. The vast repertoire of available fluorescent proteins [60] permitted us to reduce the complexity of the hardware device, avoiding moving parts such as filter wheels. Moving filters increases not only the costs but also the risk of imaging artifacts due to vibration and movement of the sample during time lapse microscopy. The genetic components were designed following advanced standards for DNA assembly that allow for cheap, rapid and combinatorial assembly of genetic systems. These assembly standards are compatible with state-or-the-art genetic libraries such as CIDAR MoClo [45] and iGEM IIs [61], permitting users access to the latest DNA components.

In this project, we declare that we have no intentions to exert protection over the genetic resources used and aim to distribute them under an open material transfer agreement that is currently being drafted [62]. Free and open source wetware has just started to gain traction (e.g. OS pharma [63], OpenPlant, [64]); and together with open access to scientific publication, data, operational procedures, open hardware and FOSS, will make science and technology more accessible, reproducible, cost-effective and equitable [51, 65–67].

Building hardware, software and genetics from scratch leads to a better understanding of technologies and their integration. New educational modalities based on “tinkering” with biology, hardware and software for STEM training and capacity building have recently emerged (e.g. [68–72]. We provide examples of simple experimental setups that can be used in practical work involving the analysis of multiple gene expression, cell-cell signaling and microbial growth. We thus hope that this project will contribute to the democratization of science, technology and innovation in education.

## Acknowledgments

We would like to thank Toby Wenzel for guidance on Docubricks documentation, Tom Baden for feedback and advice on camera, Bernardo Pollak for helping with sequences. Douglas Densmore for the CIDAR MoClo Parts Kit. We would like to thank reviewers for very valuable comments.

OpenPlant Fund for financial support to INN, TFM, RHP, JEK, TCM, TJR, and FF, Fondecyt Iniciacion 11140776 and 11161046 for providing financial support to FF and TJR, respectively. Fondo de Desarrollo de Areas Prioritarias (FONDAP) Center for Genome Regulation (15090007) and Millennium Nucleus Center for Plant Systems and Synthetic Biology (NC130030) for providing financial support to FF. Fondecyt Regular1150430 for providing financial support to JEK. "The funders had no role in study design, data collection and analysis, decision to publish, or preparation of the manuscript."

## Supporting Information

**S1 Figure: sfGFP expression by promoters of different strength. S2 Figure: Image of DNA electrophoresis gel.**

**S1 Appendix. Assembly instructions for top and bottom enclosures S2 Appendix. Assembly instructions for Raspberry Pi camera**

**S3 Appendix. Assembly instructions for camera system S4 Appendix. assembly instructions for PCB**

**S5 Appendix. Fluorescent signal mathematical analysis for time lapse images S6 Appendix. Colony classification maths**

**S1 Movie. Time lapse of diffusion assay for E. coli cells producing (“sender cells”) and responding to (“receiver cells”) C6 and C12 homoserine lactones plated on permeable membranes printed with hydrophobics grid lines.**

**S2 Movie. Time lapse of “star” diffusion assay for E. coli cells producing (“sender cells”) and responding to (“receiver cells”) C6 and C12 homoserine lactones plated on permeable membranes printed with hydrophobics grid lines (compressed for web streaming).**

**S3 Movie. Time lapse of sectoring colony growth (compressed for web streaming).**

**S4 Movie. Time lapse of mixed fluorescent bacteria (compressed for web streaming).**

**S5 Movie. Time lapse of growing E. coli labelled with sfGFP (compressed for web streaming)**

**S6 Movie. Time lapse of growing E. coli labelled with CyOFP. S7 Movie. Time lapse of growing E. coli labelled with mBeRFP.**

**S8 Movie. Time lapse of mixed E. coli labelled with mBeRFP, sfGFP and mBeRFP.**

**S1 Figure:**
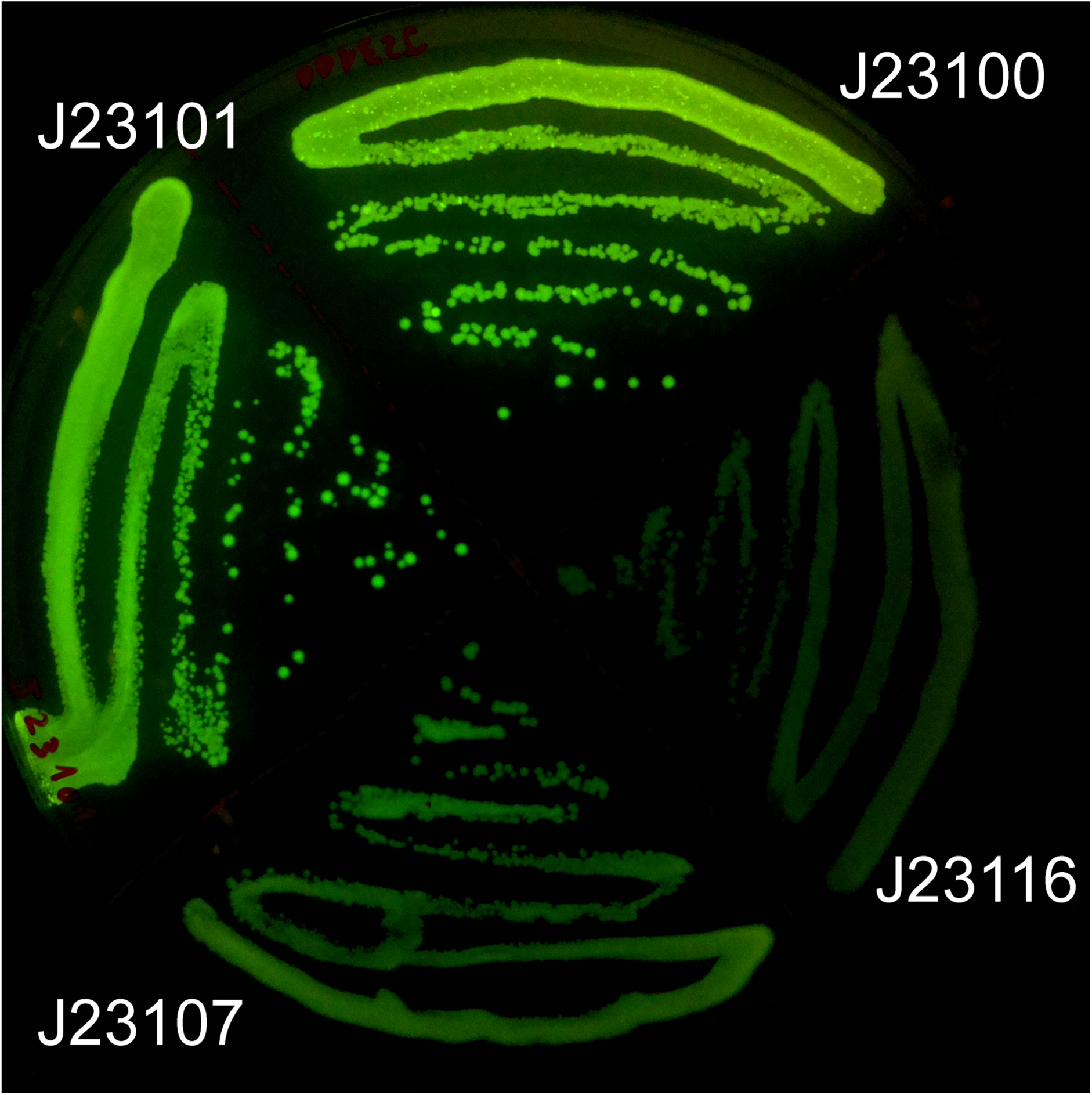
sfGFP expression by promoters of different strength: 1: J23100, 2: J23101, 3: J23107 and 4: J23116. BCD12 RBS and B0015 terminator was used for all the combinations.

**S2 Figure:**
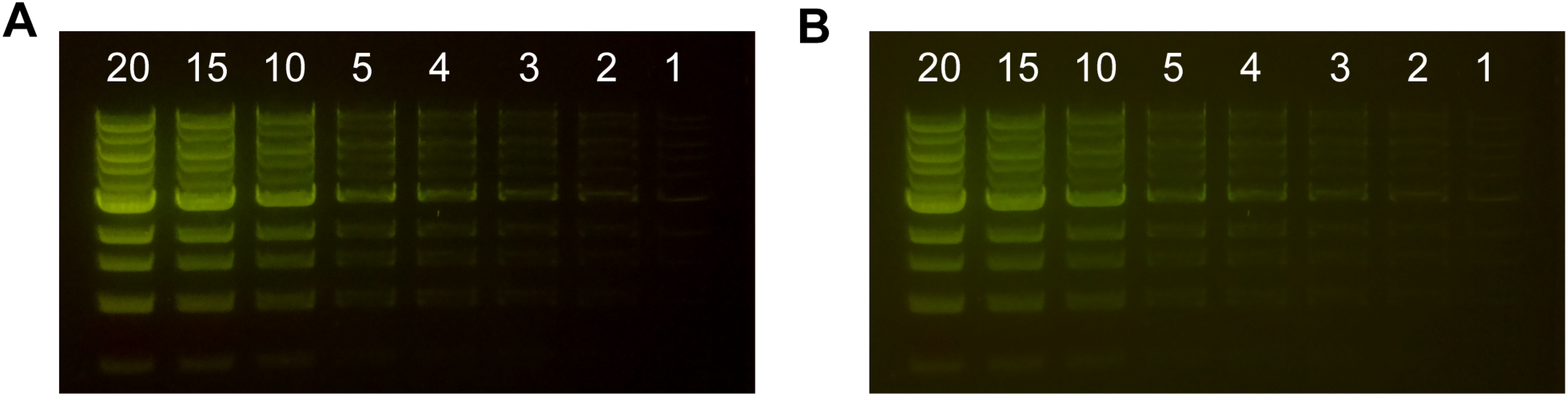
Image of DNA electrophoresis gel. (A) Gel loaded with 20, 15, 10, 5, 4,3,2 and 1 l of 1 Kb Ladder (NEB). and imaged using raspistill command: -t 5000 -ss 240000 -ISO 300 -awbg 1,1 -co 30. (B) using raspistill command: -t 5000 -ss 170000 -ISO 300 -awbg 1,1 -co 0.

## S1 Appendix. Assembly instructions for top and bottom enclosures

Fluorescence Imaging Station

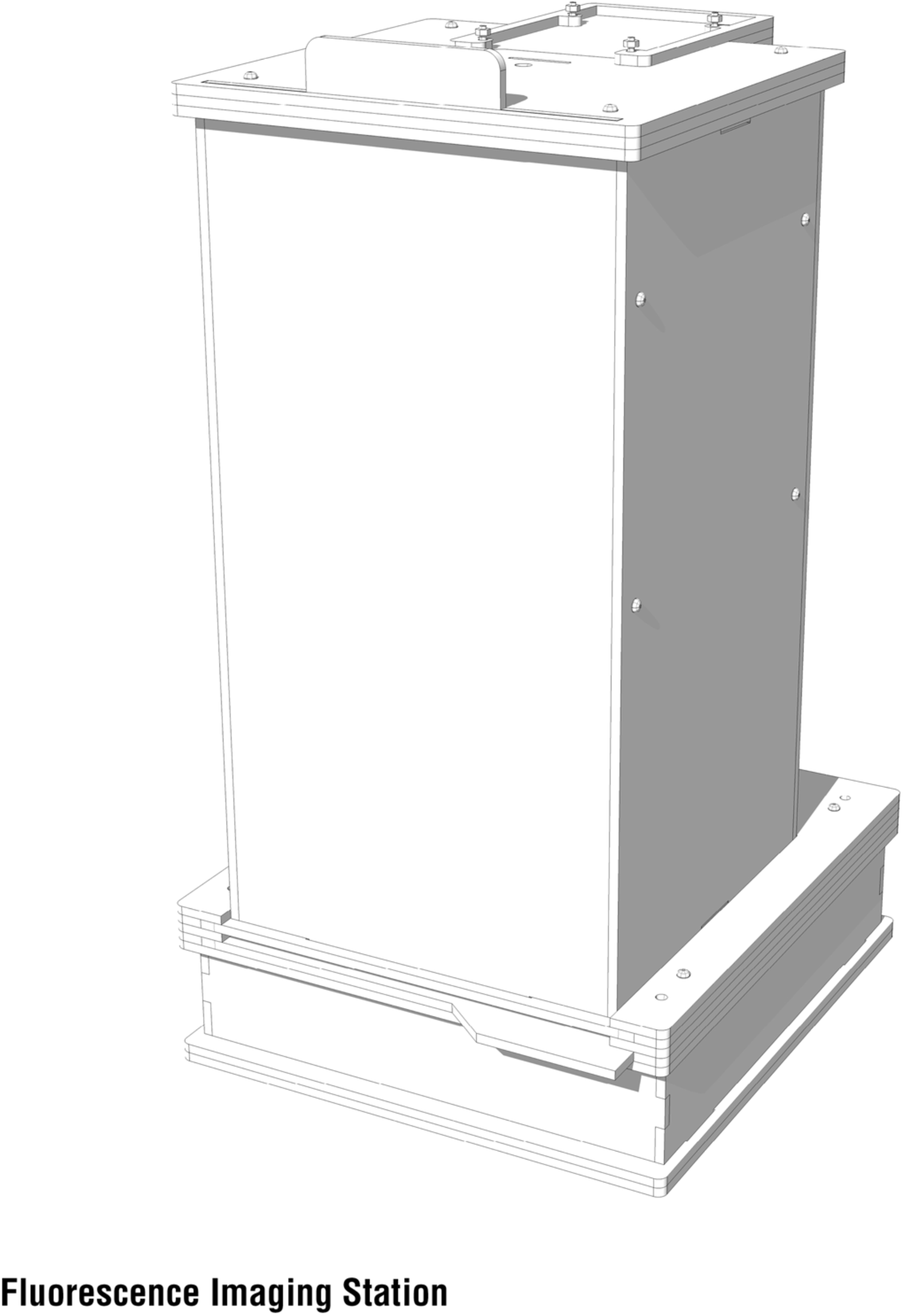

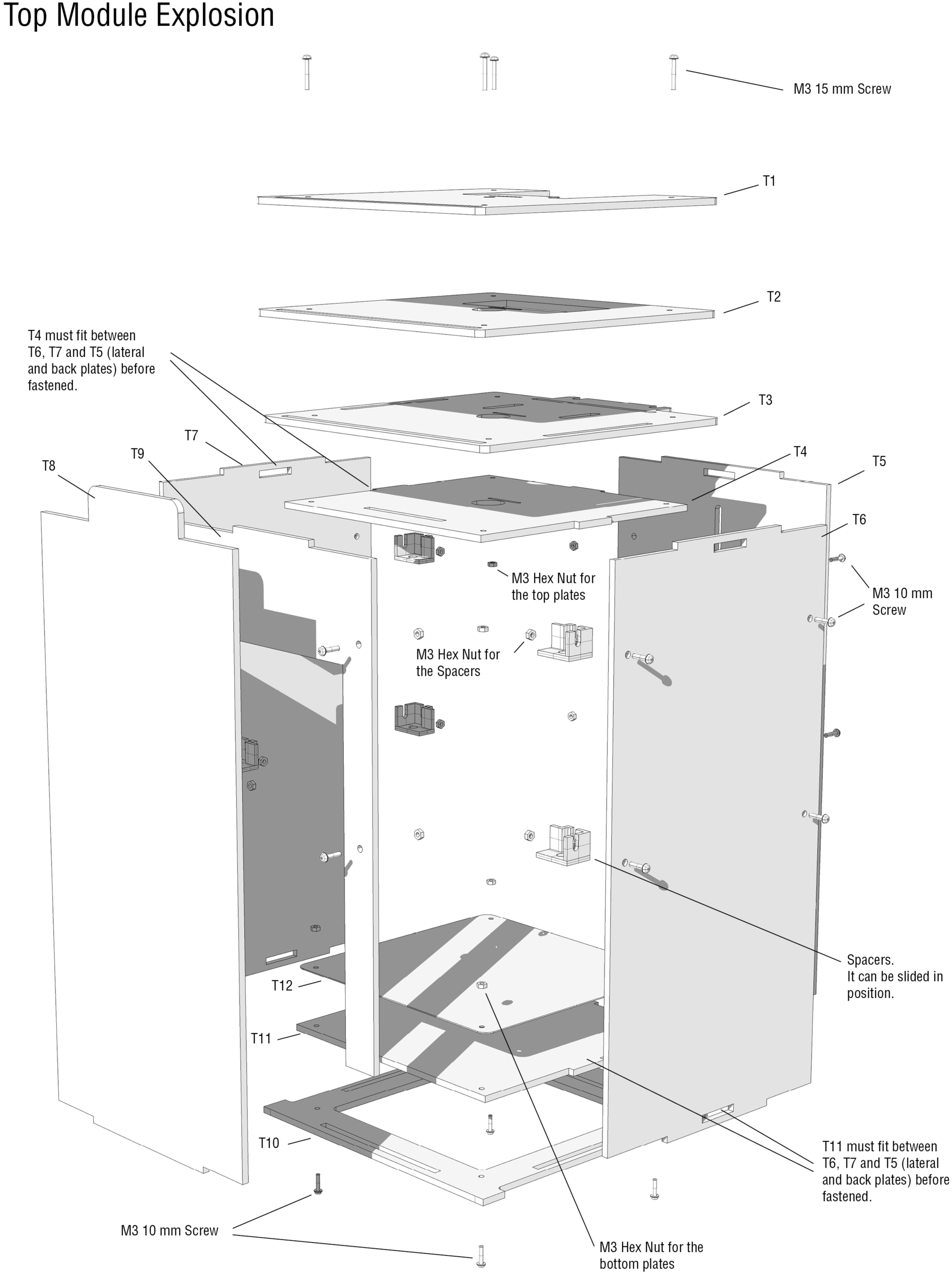

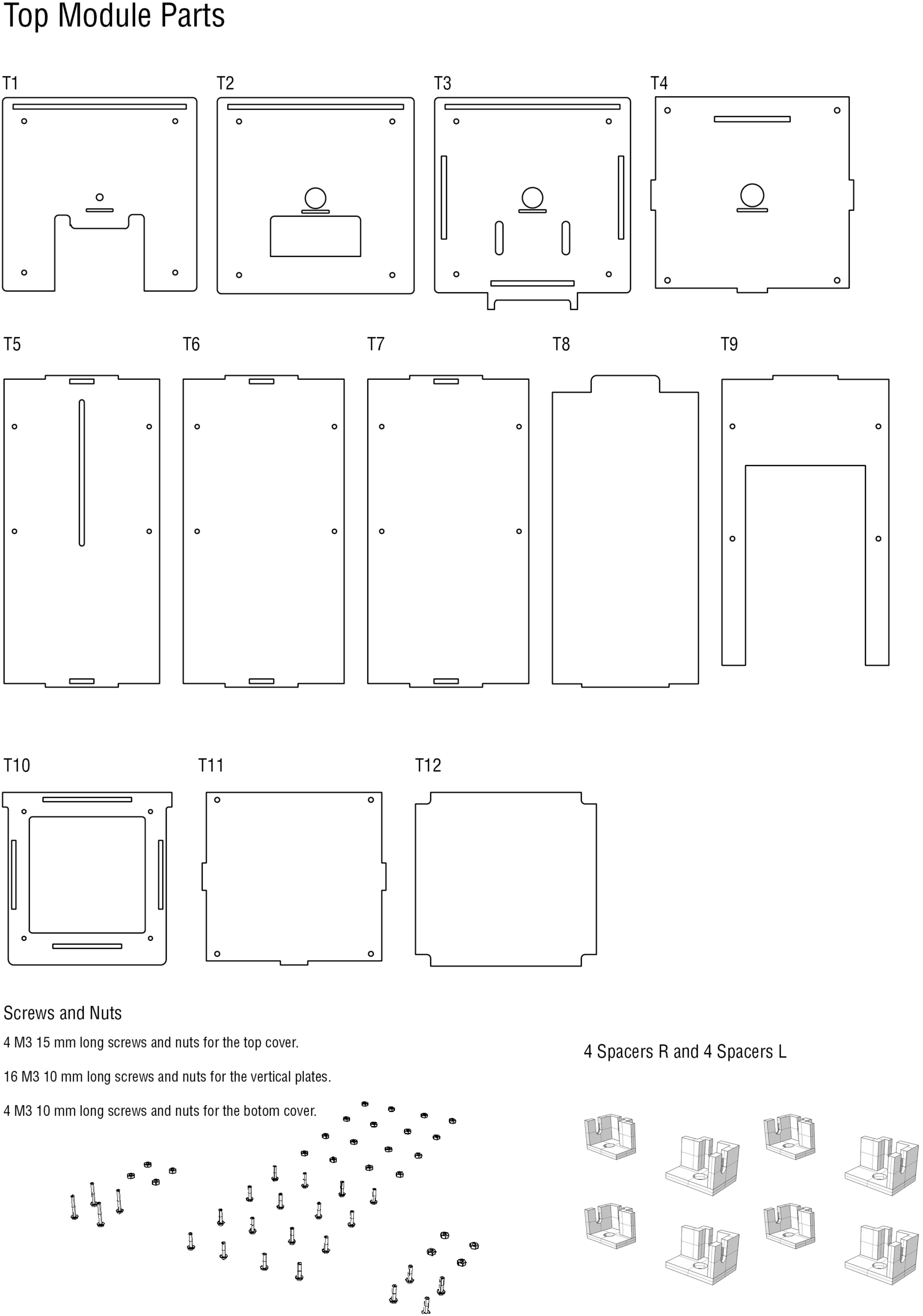

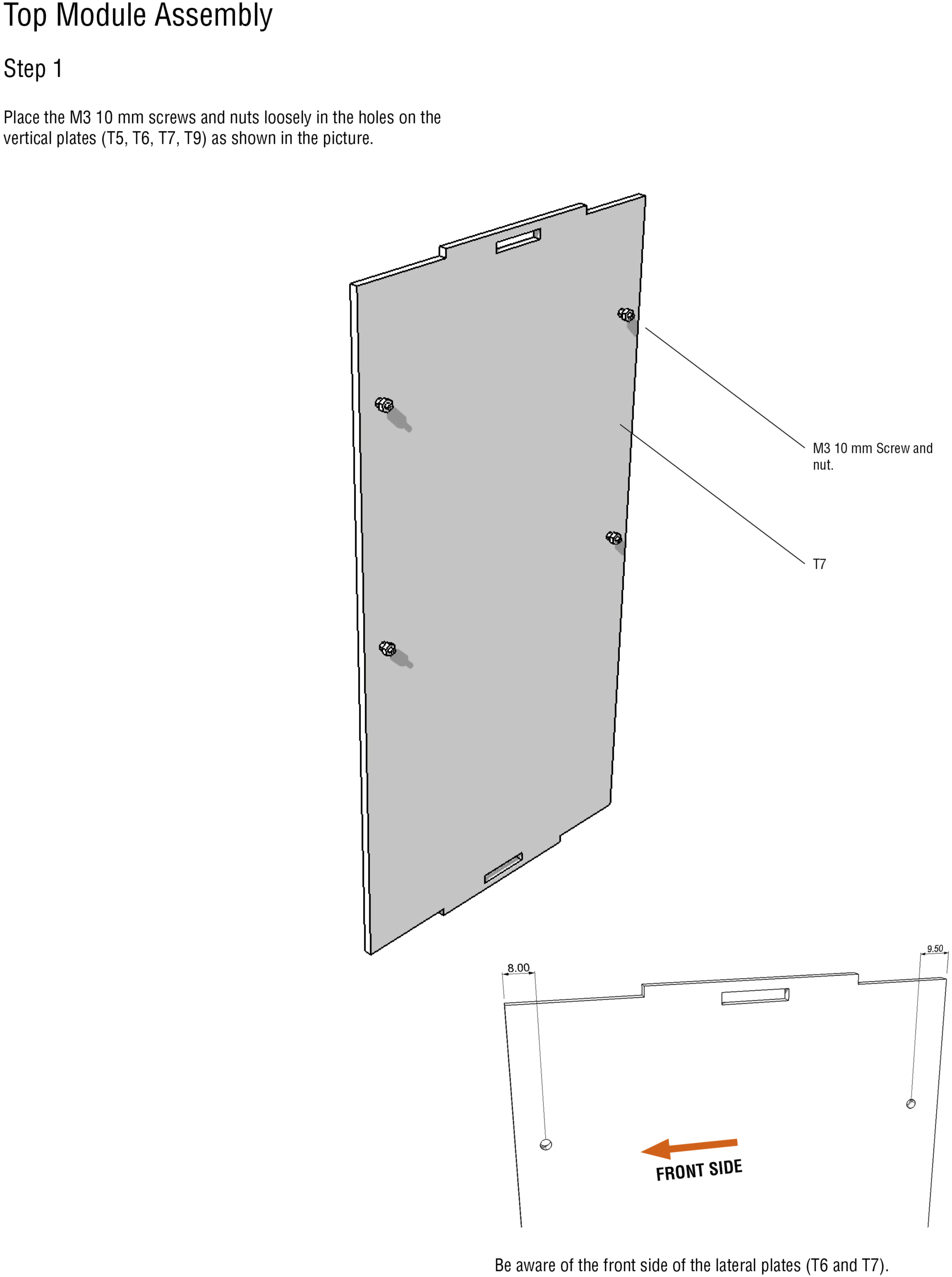

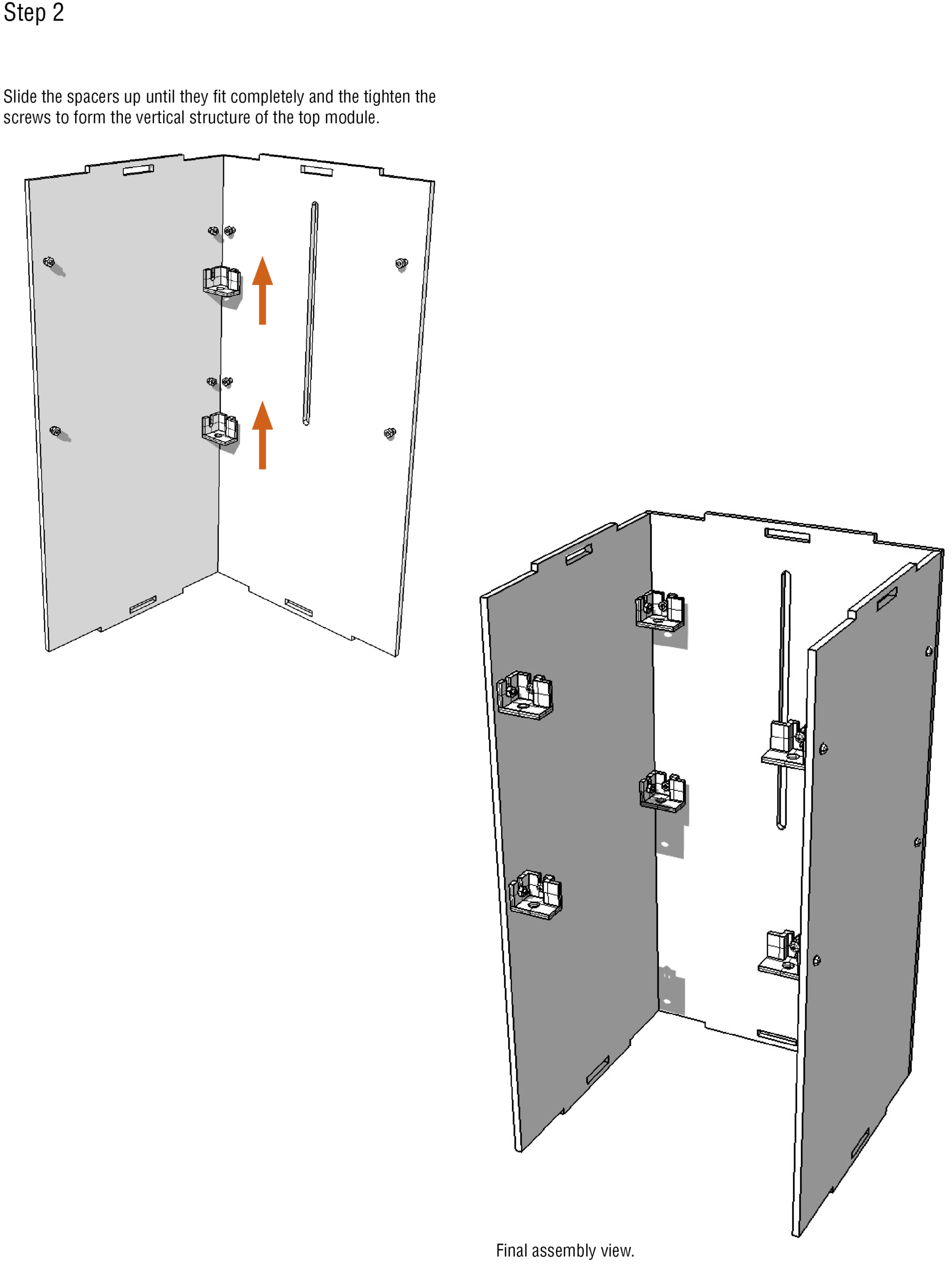

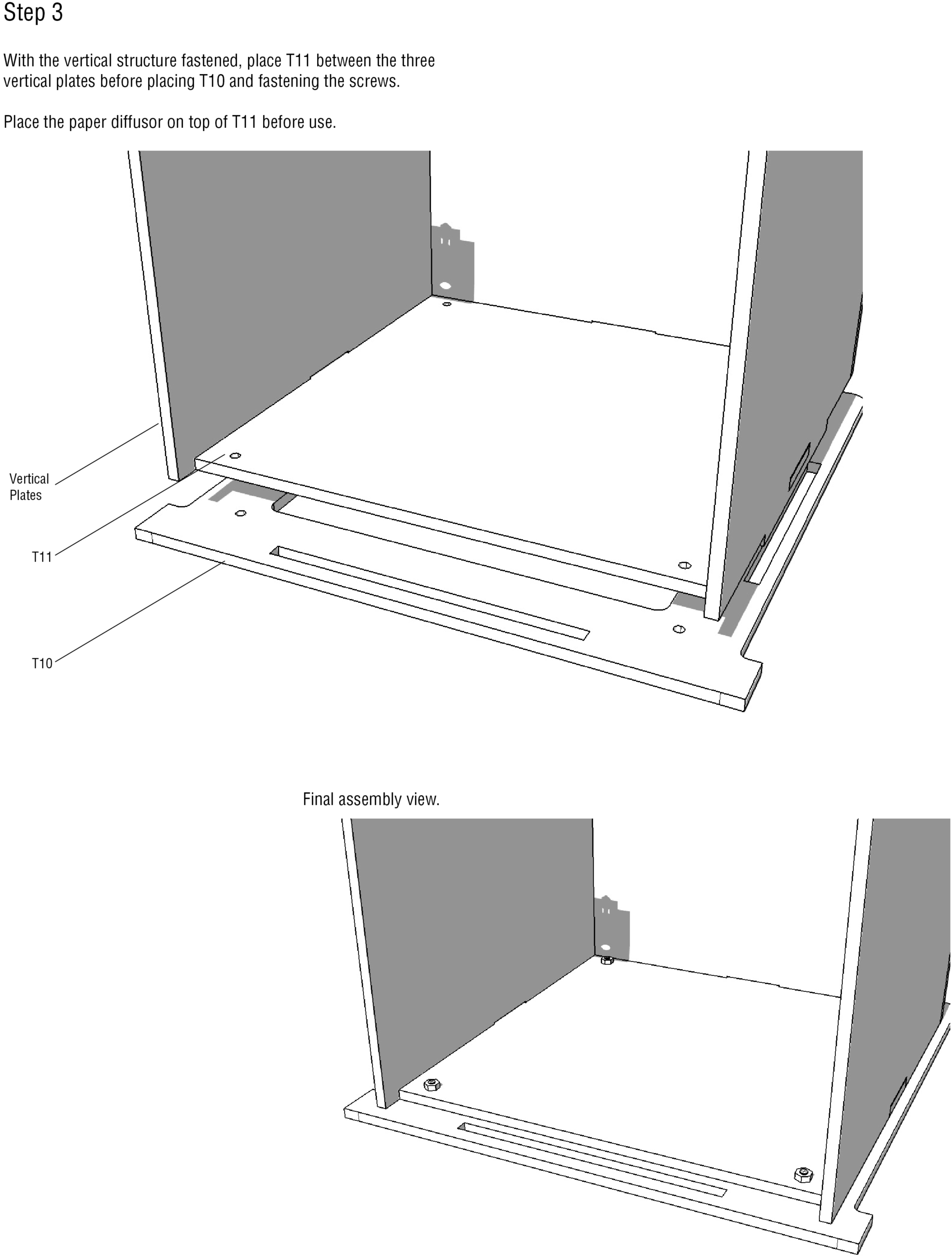

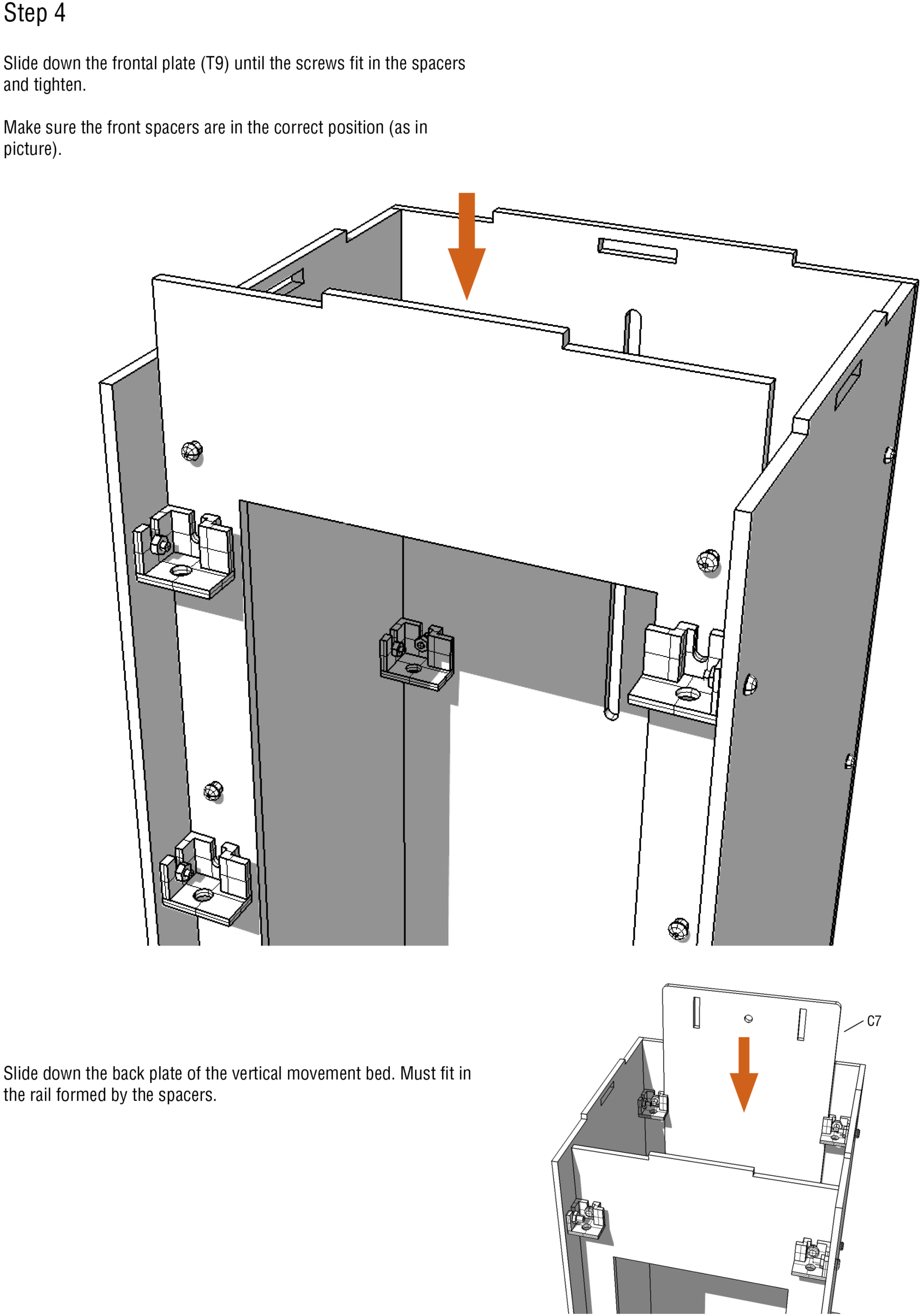

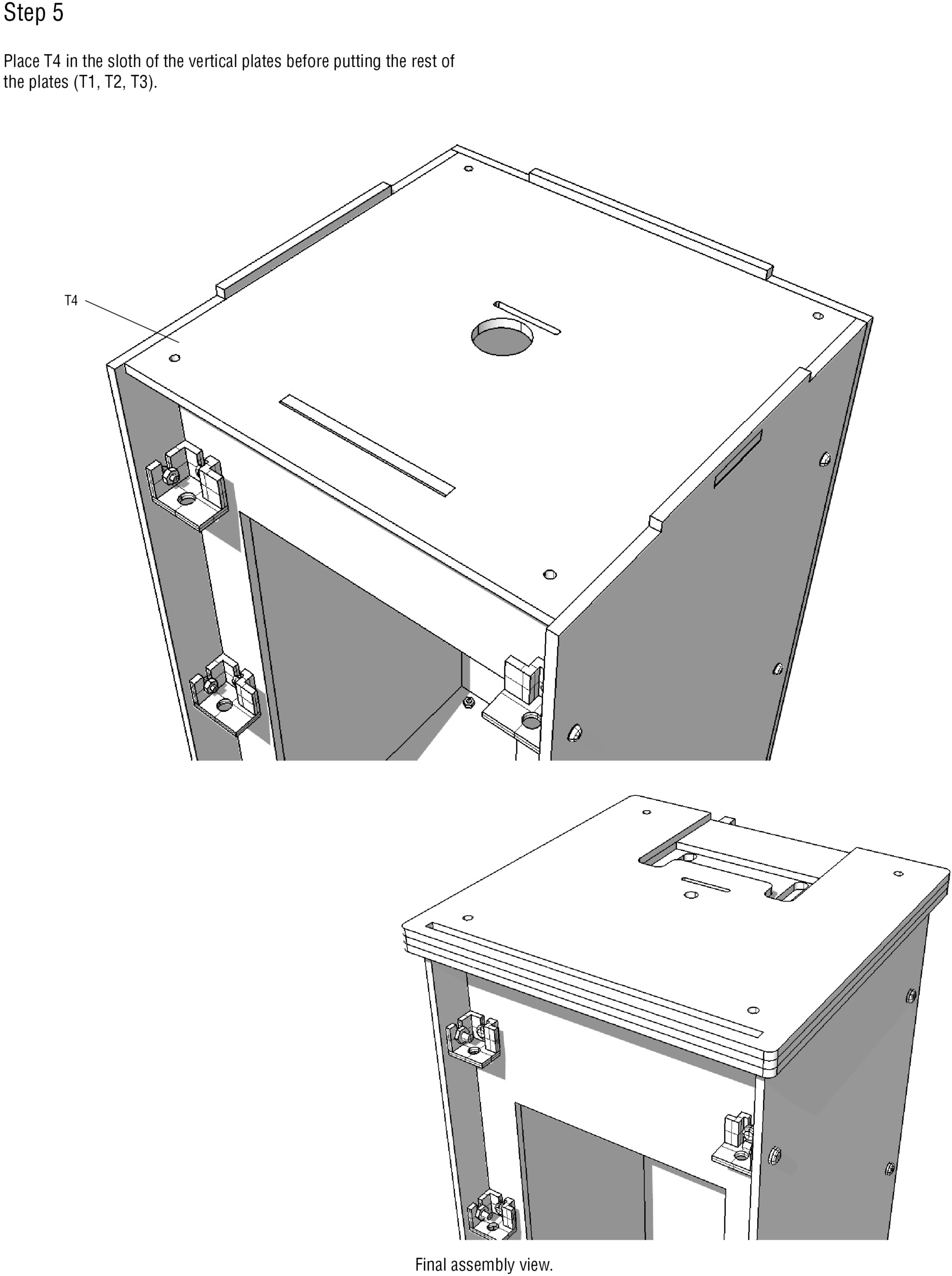

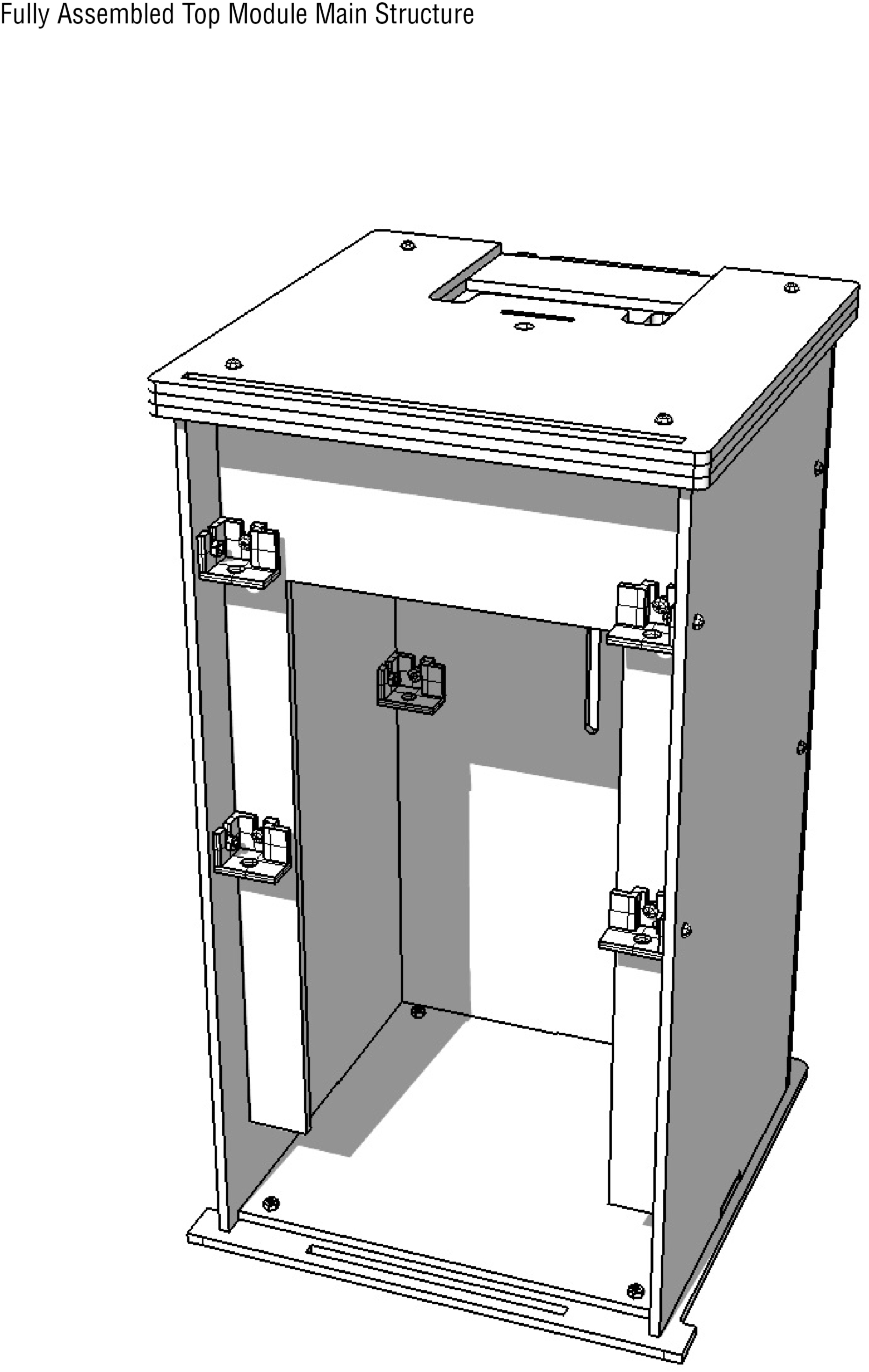

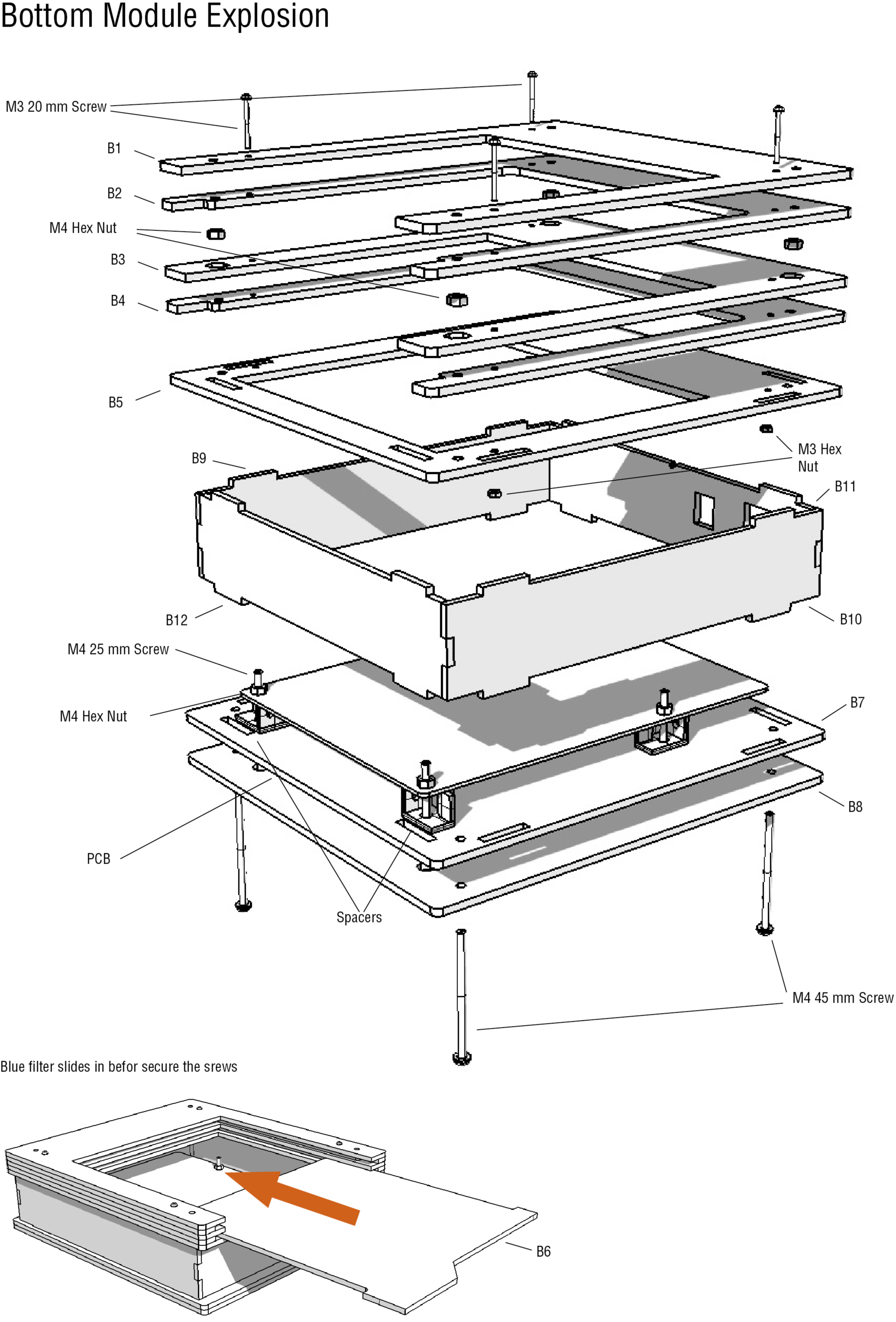

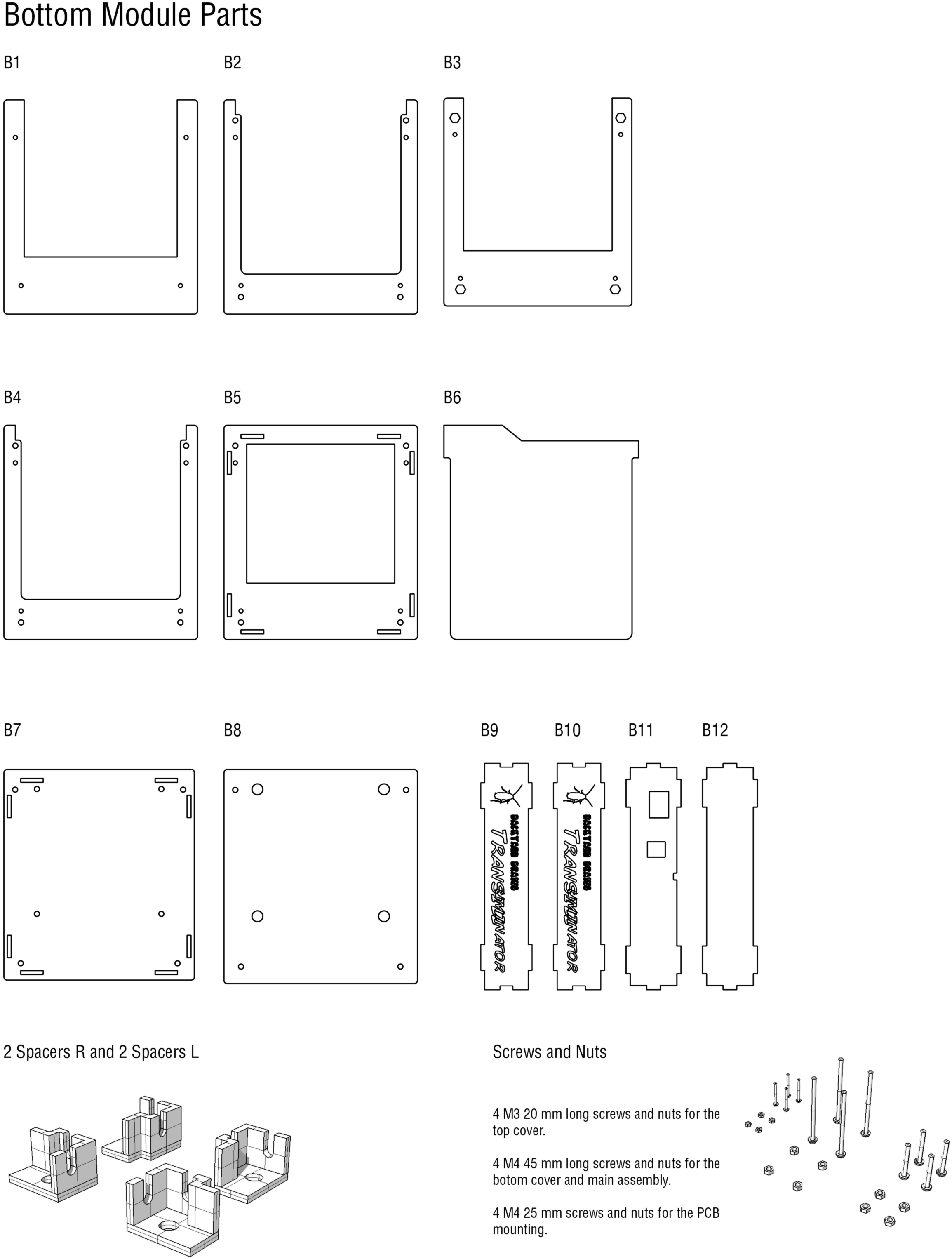

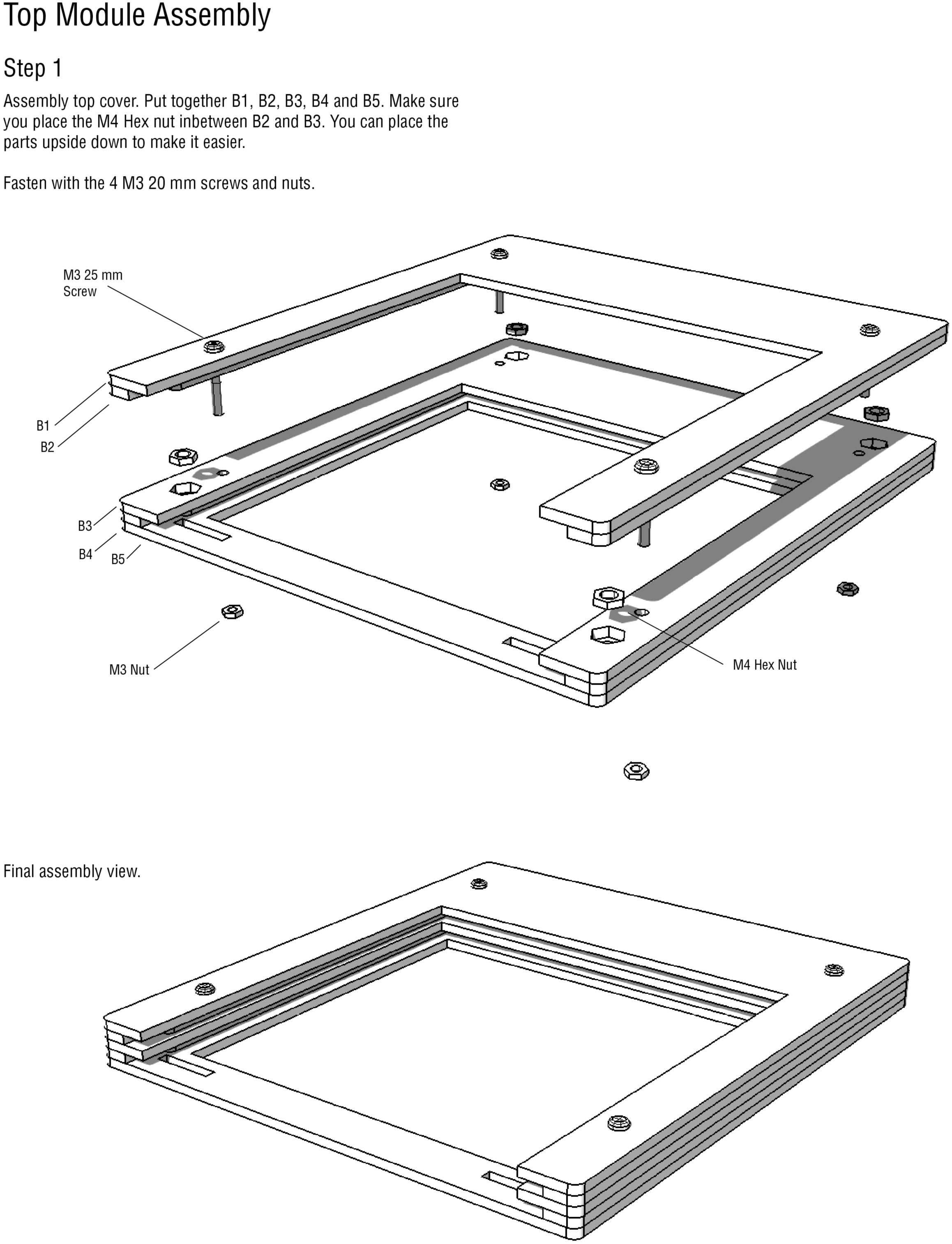

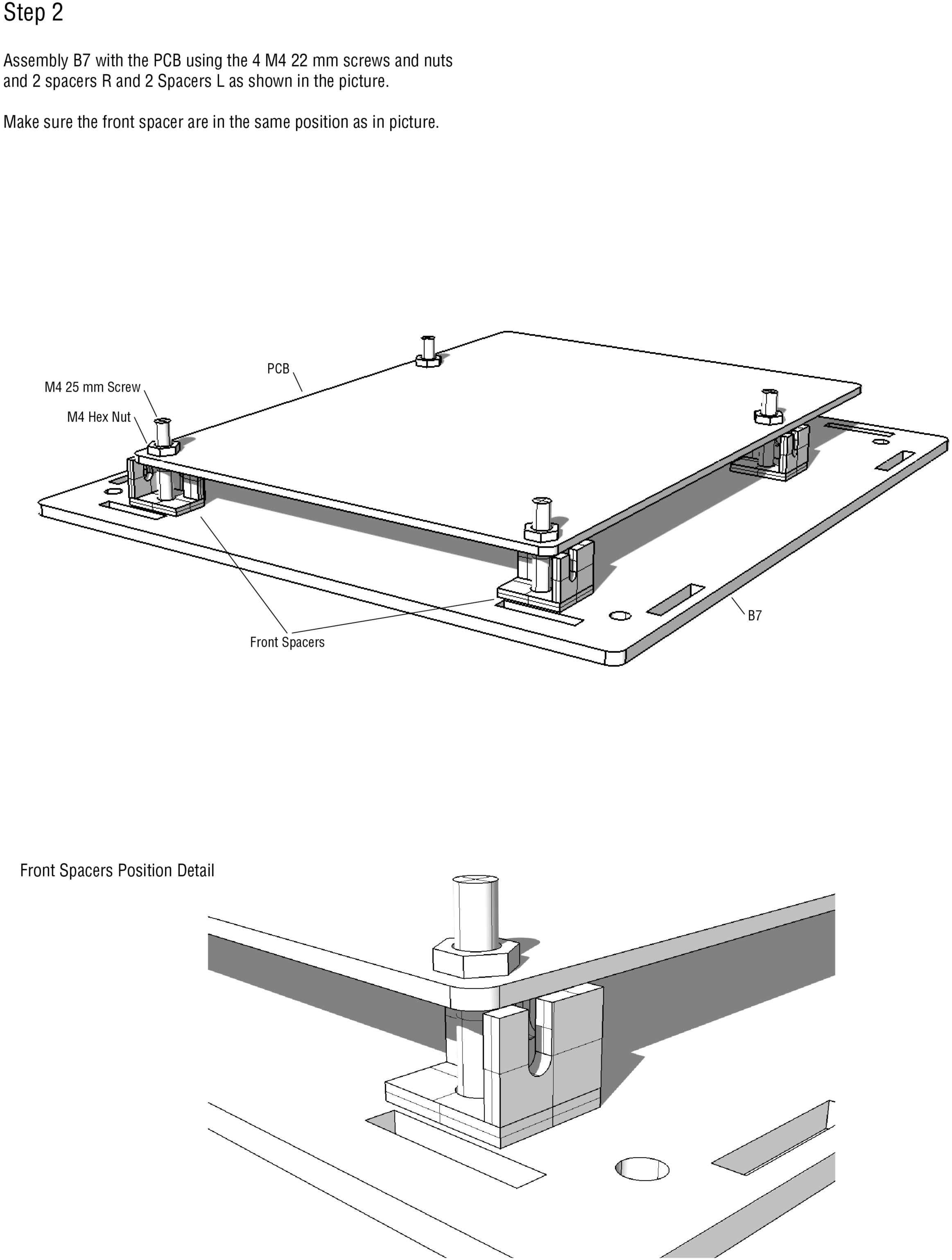

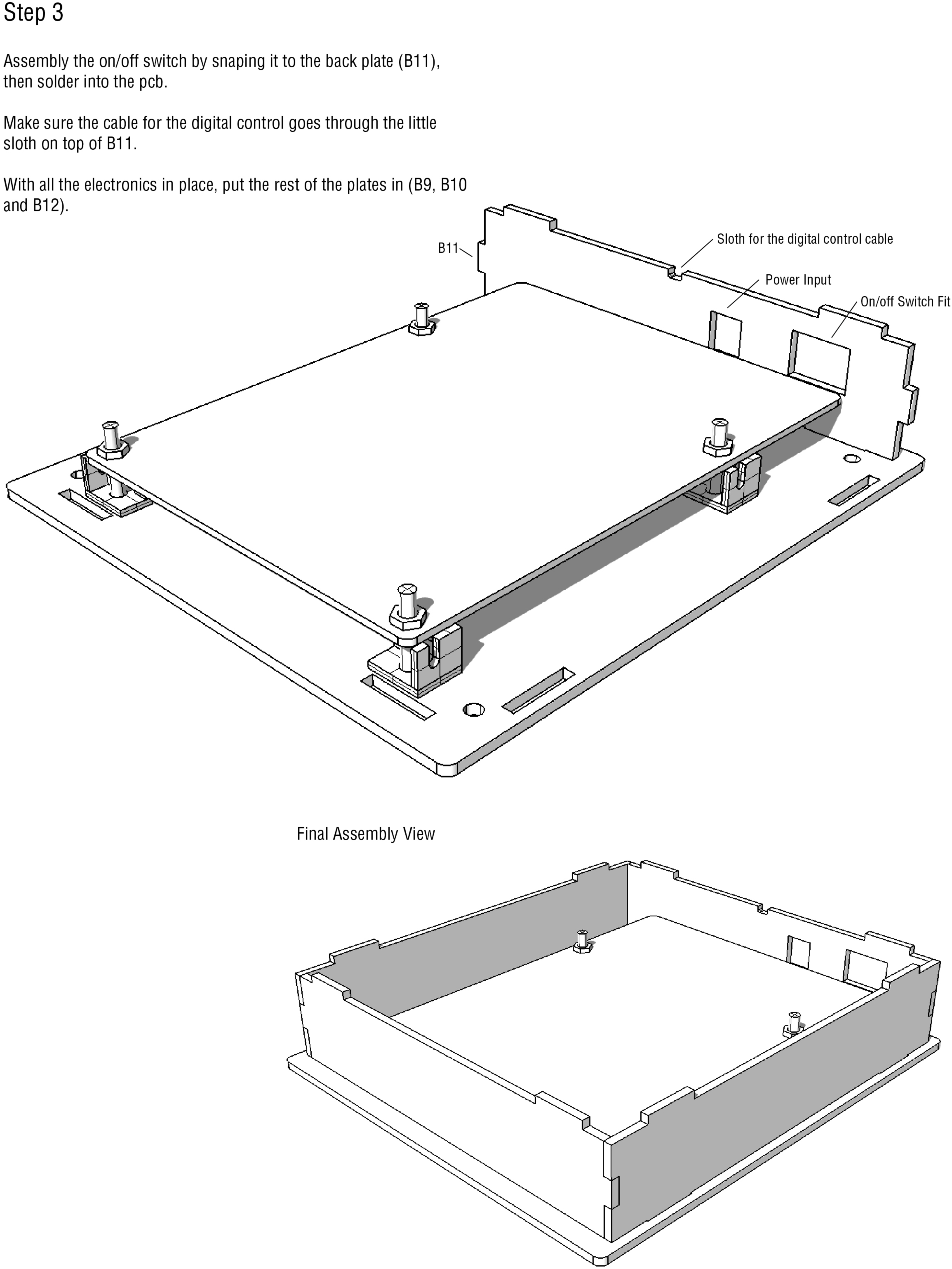

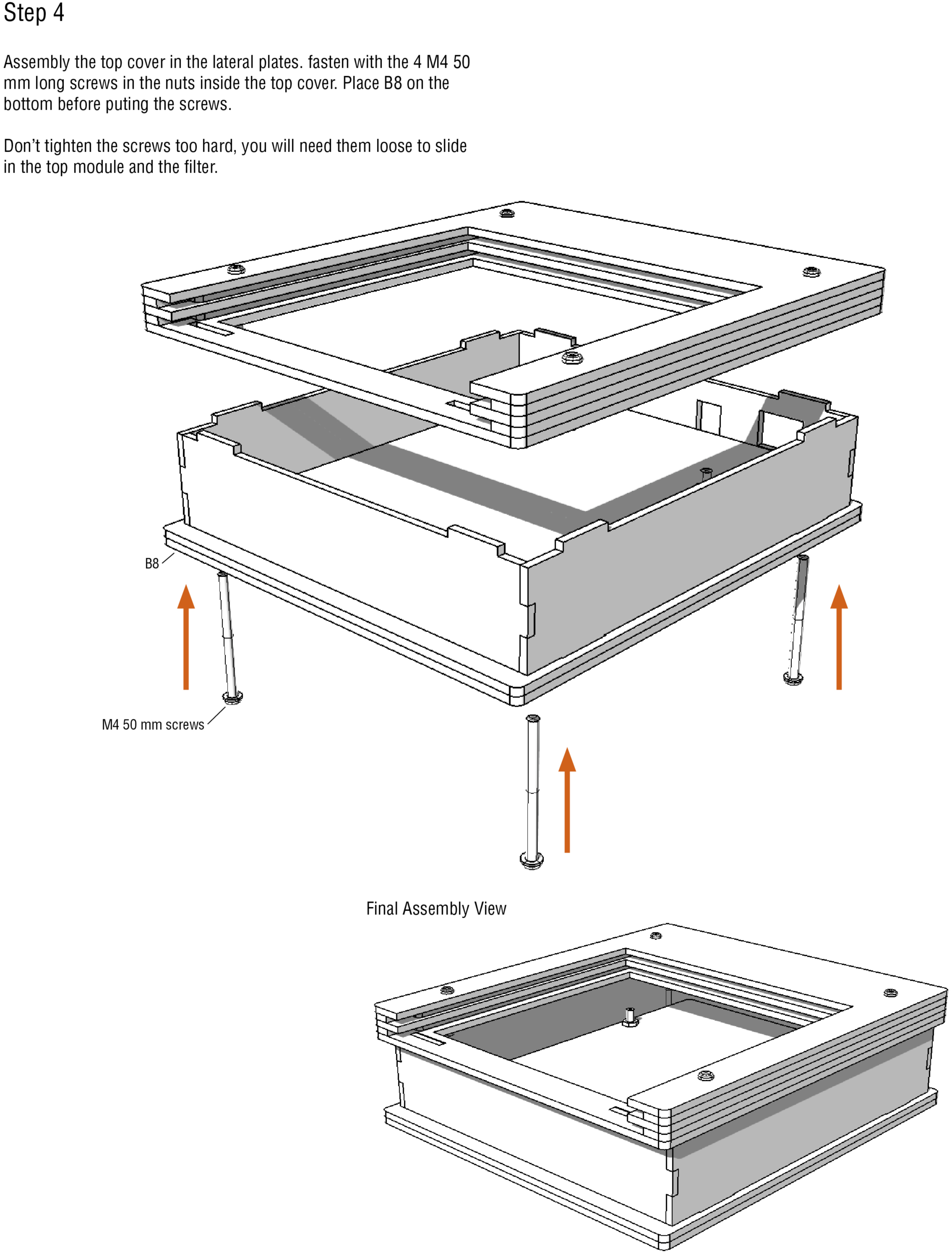

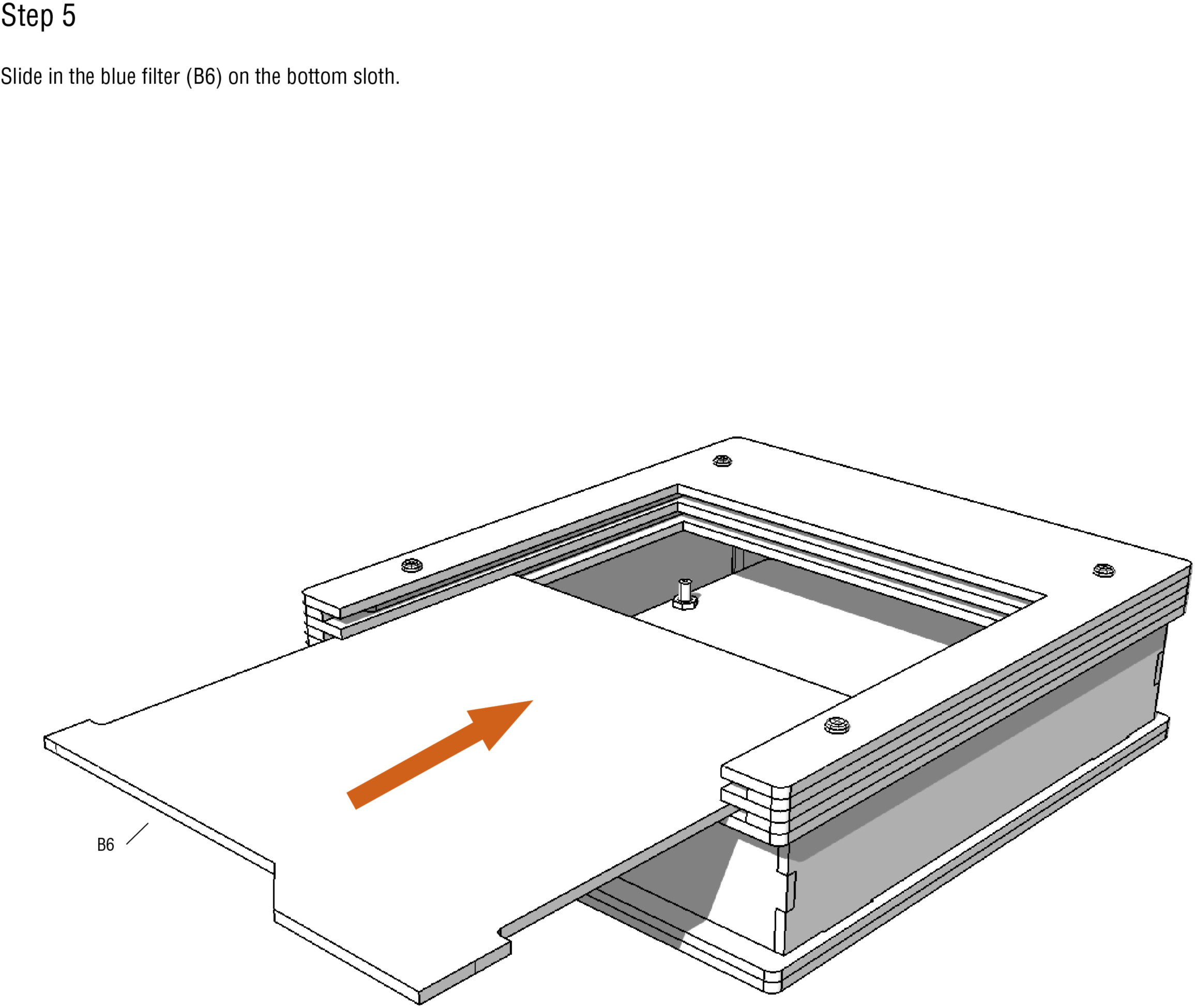

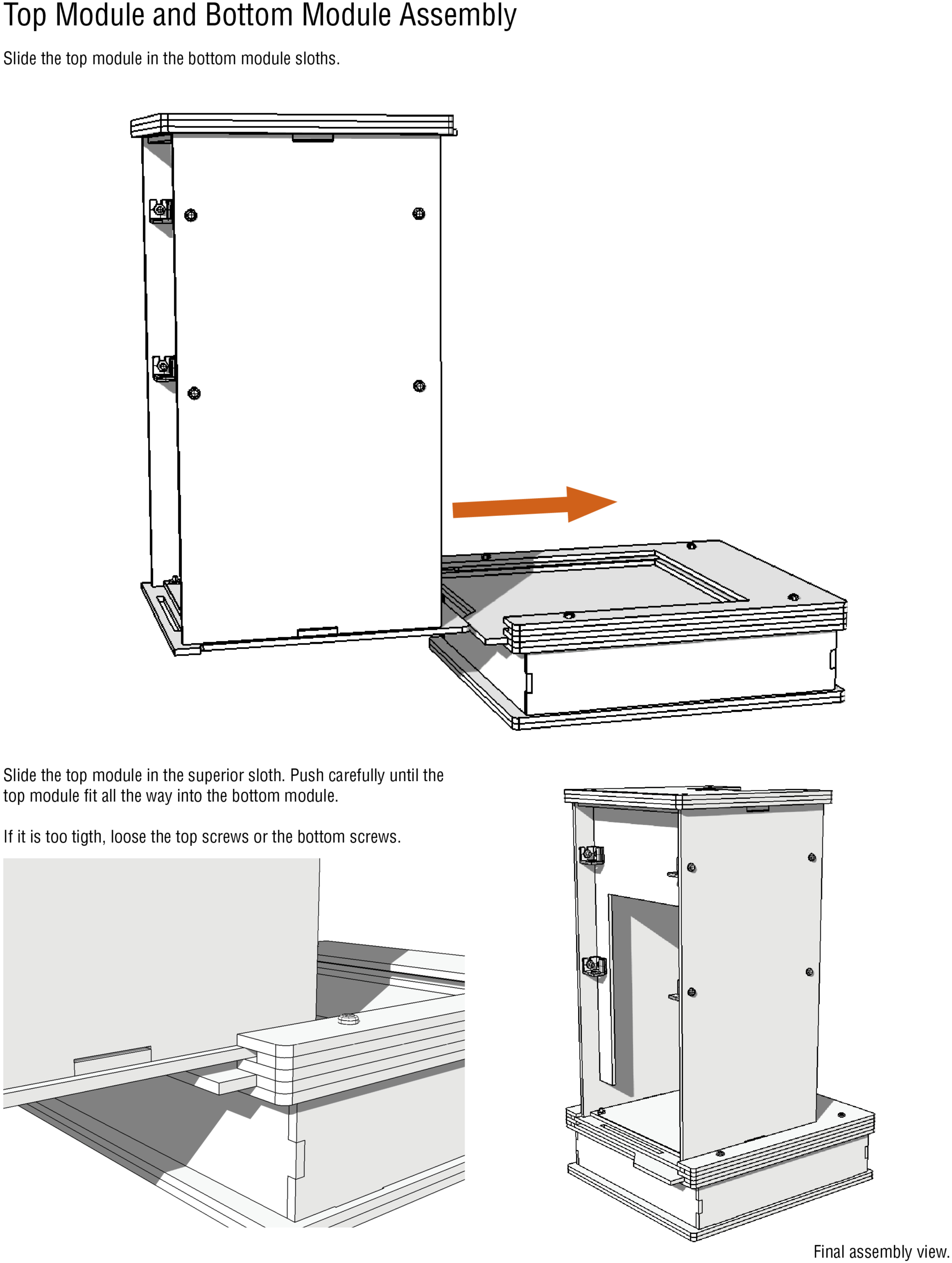

## S2 Appendix. Assembly instructions for Raspberry Pi camera

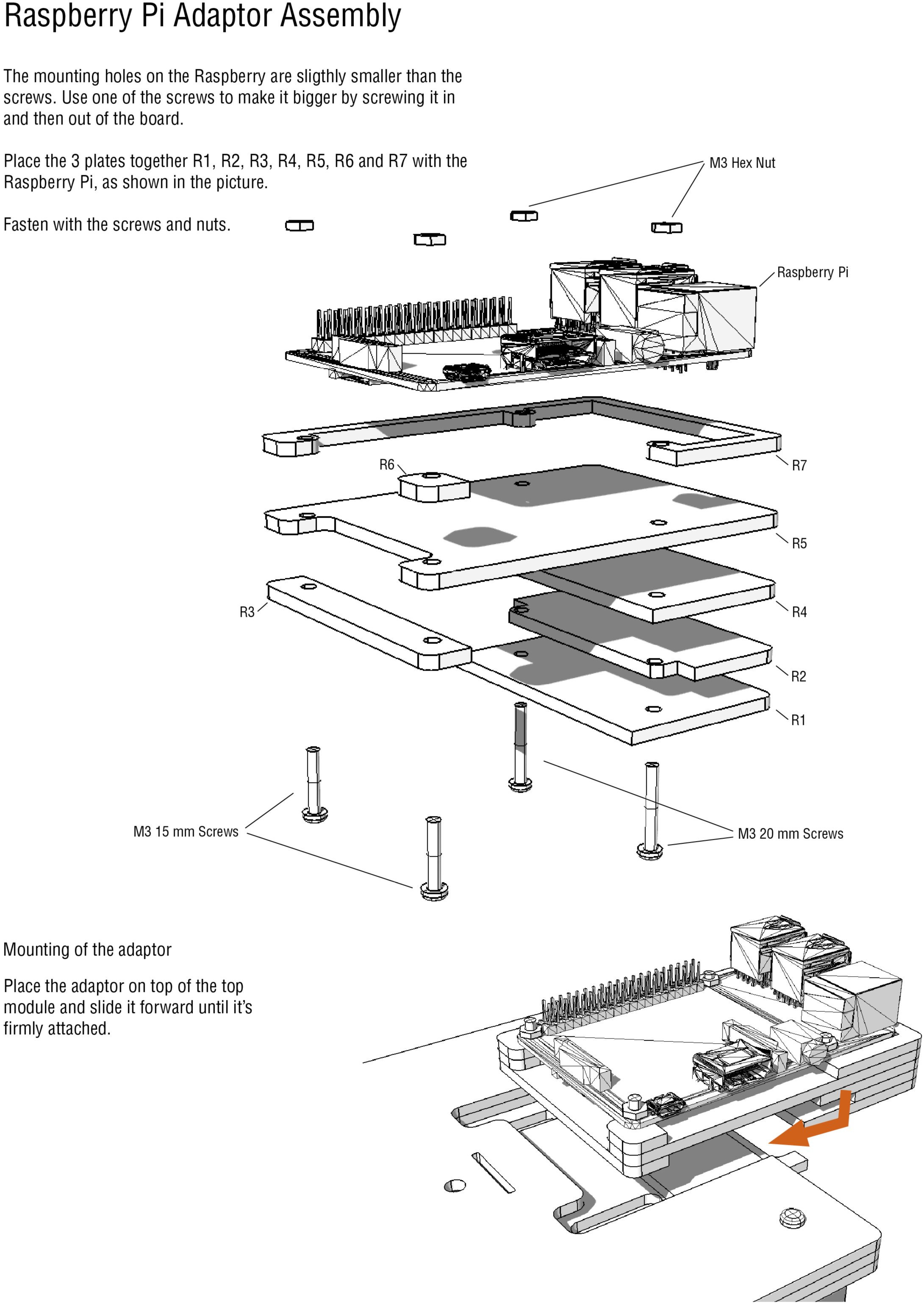

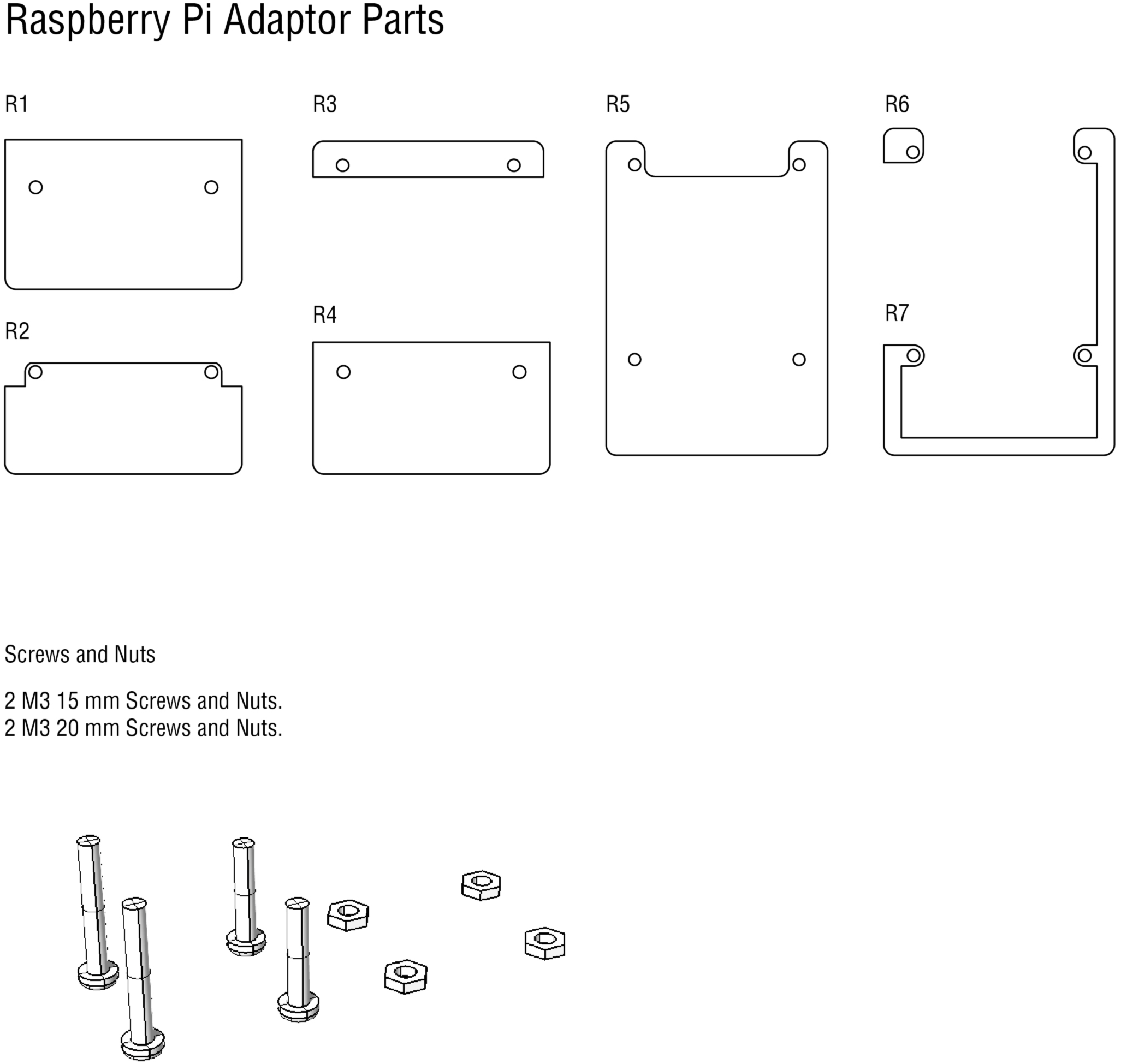

## S3 Appendix. Assembly instructions for camera system

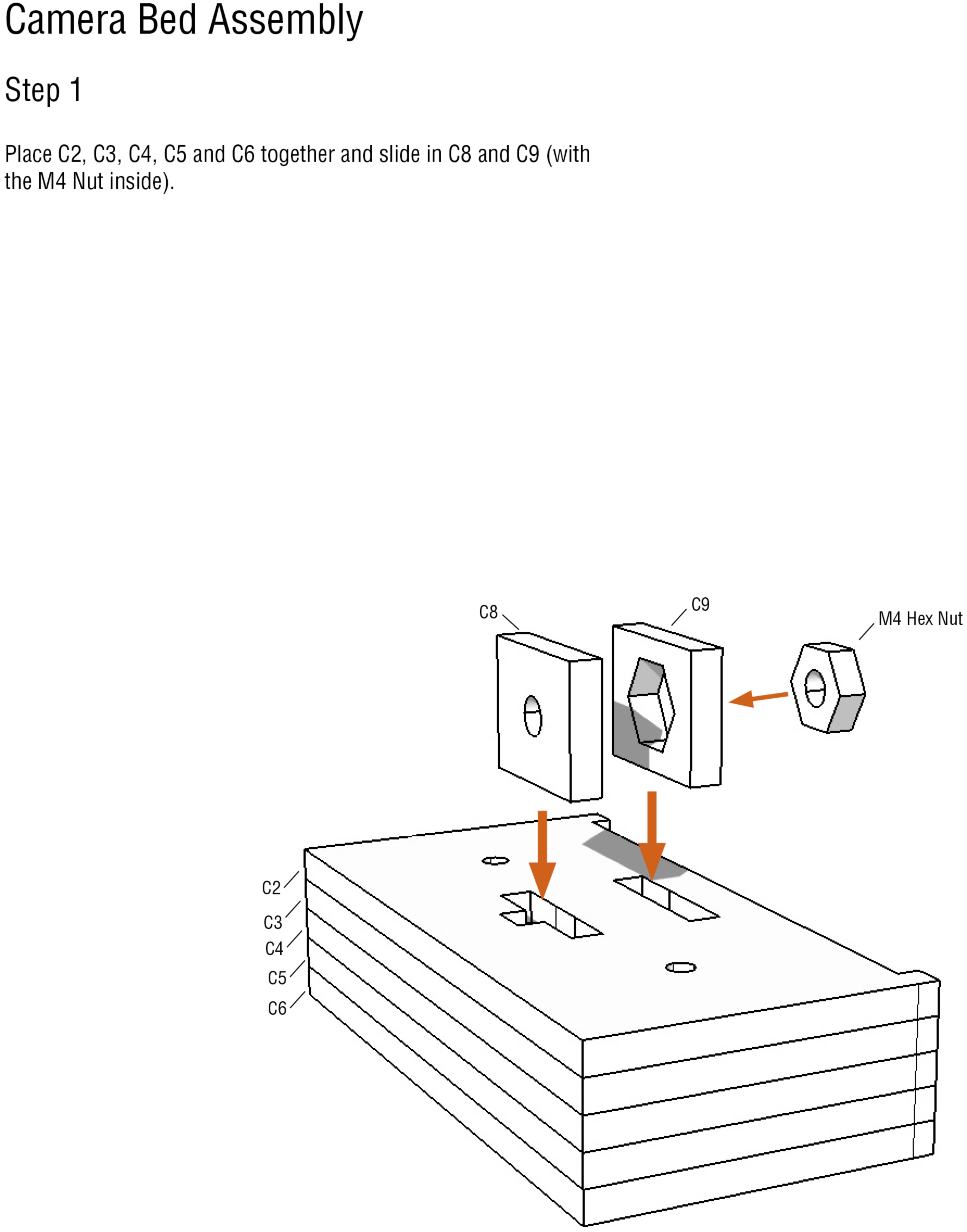

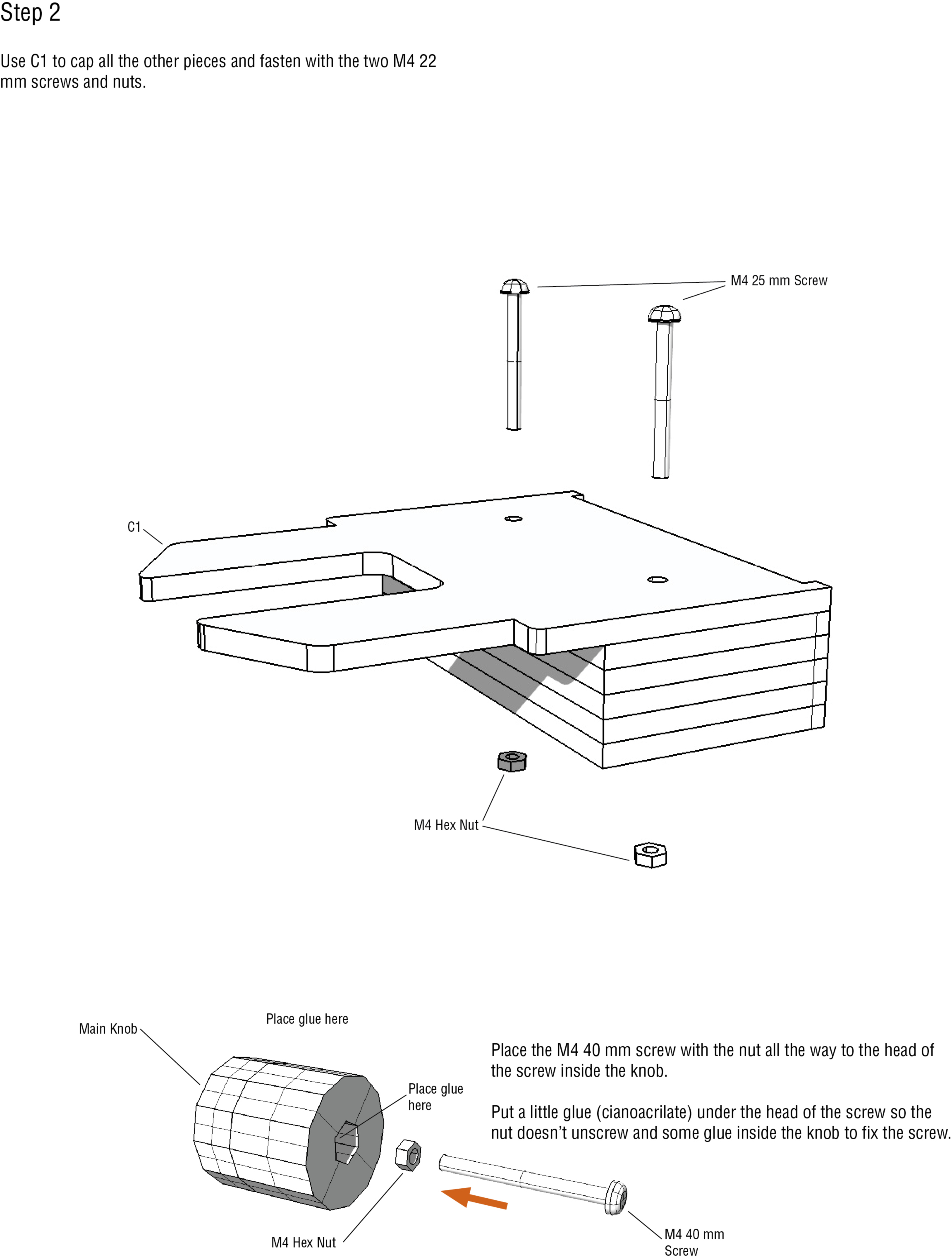

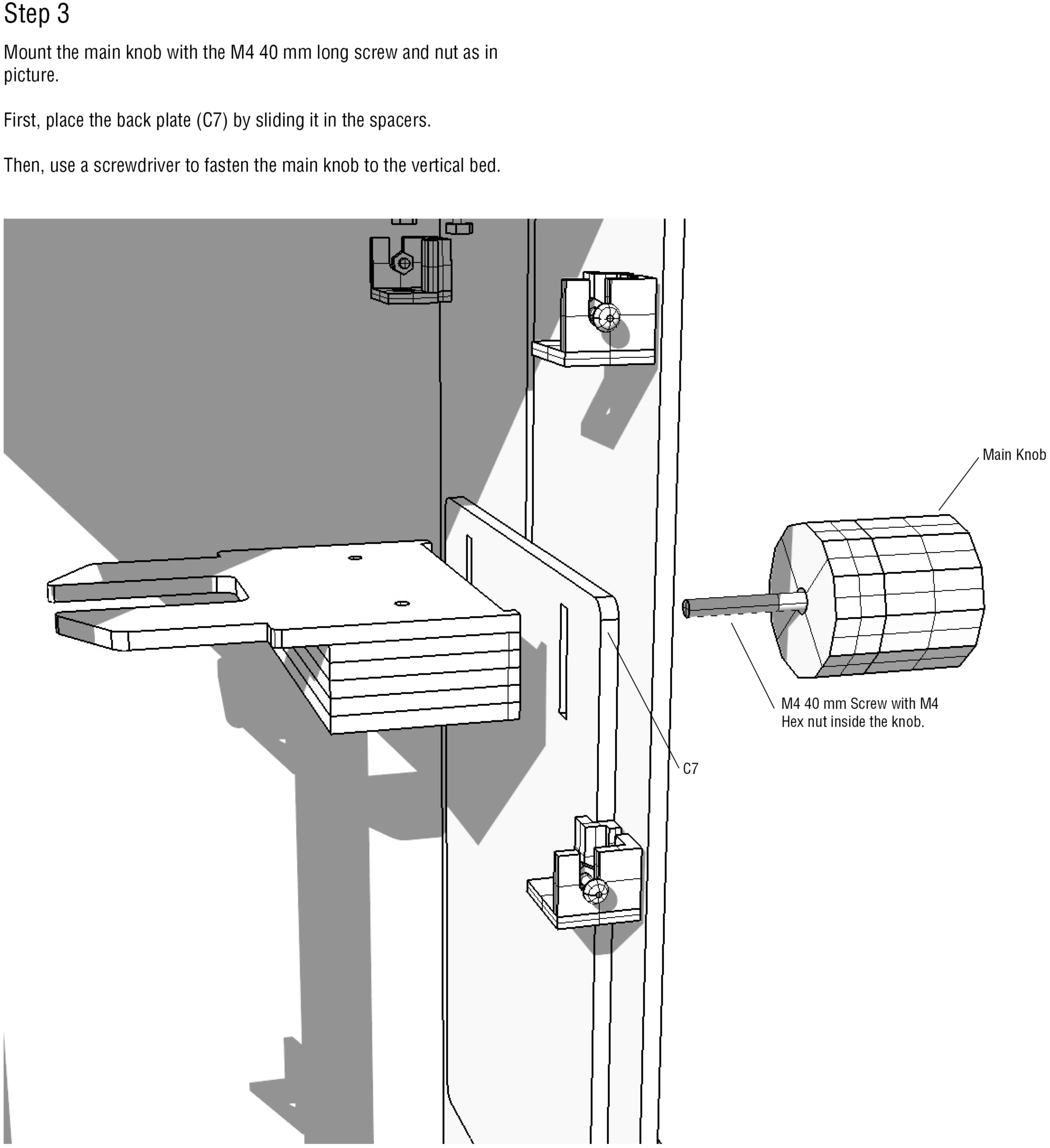

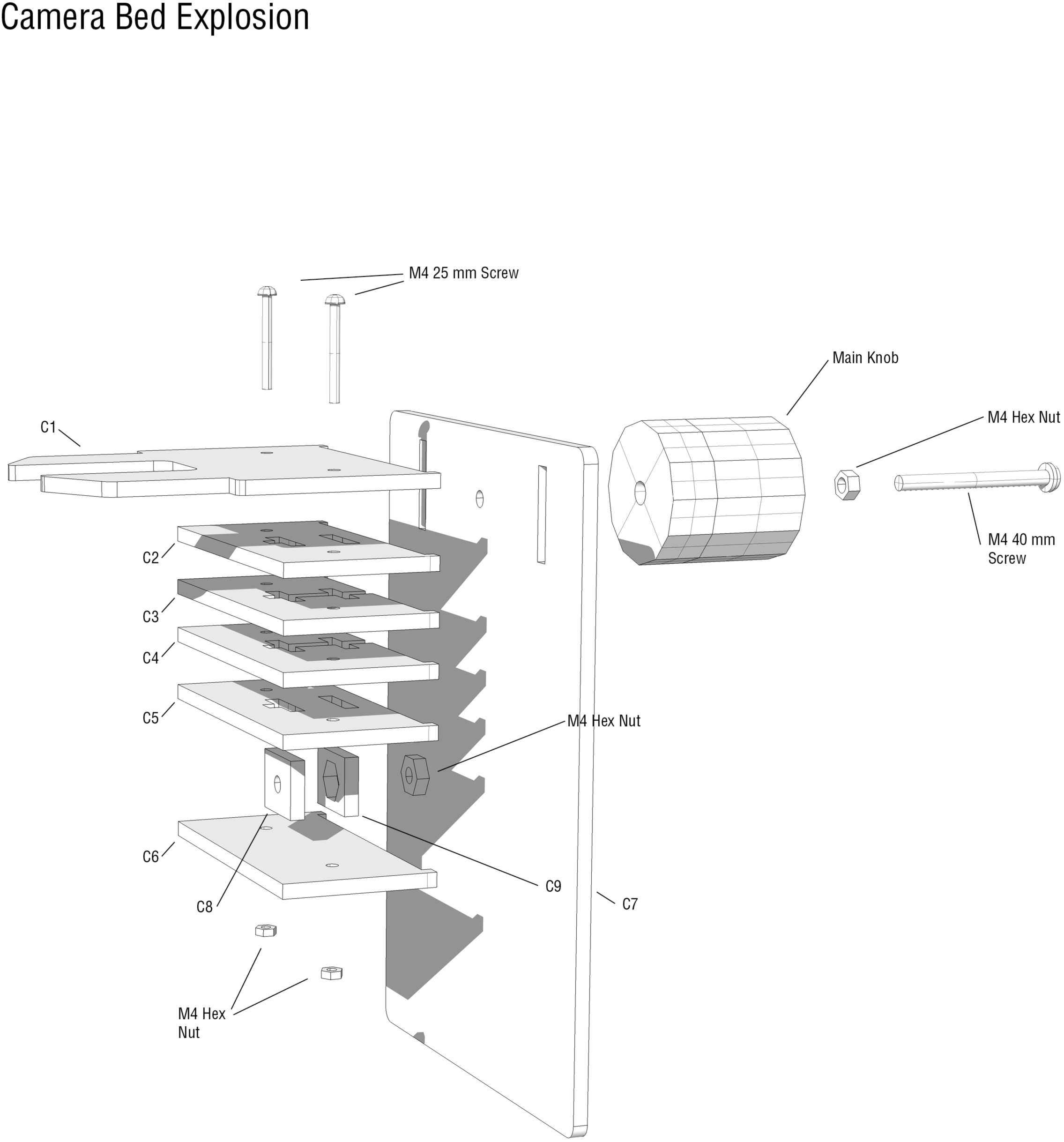

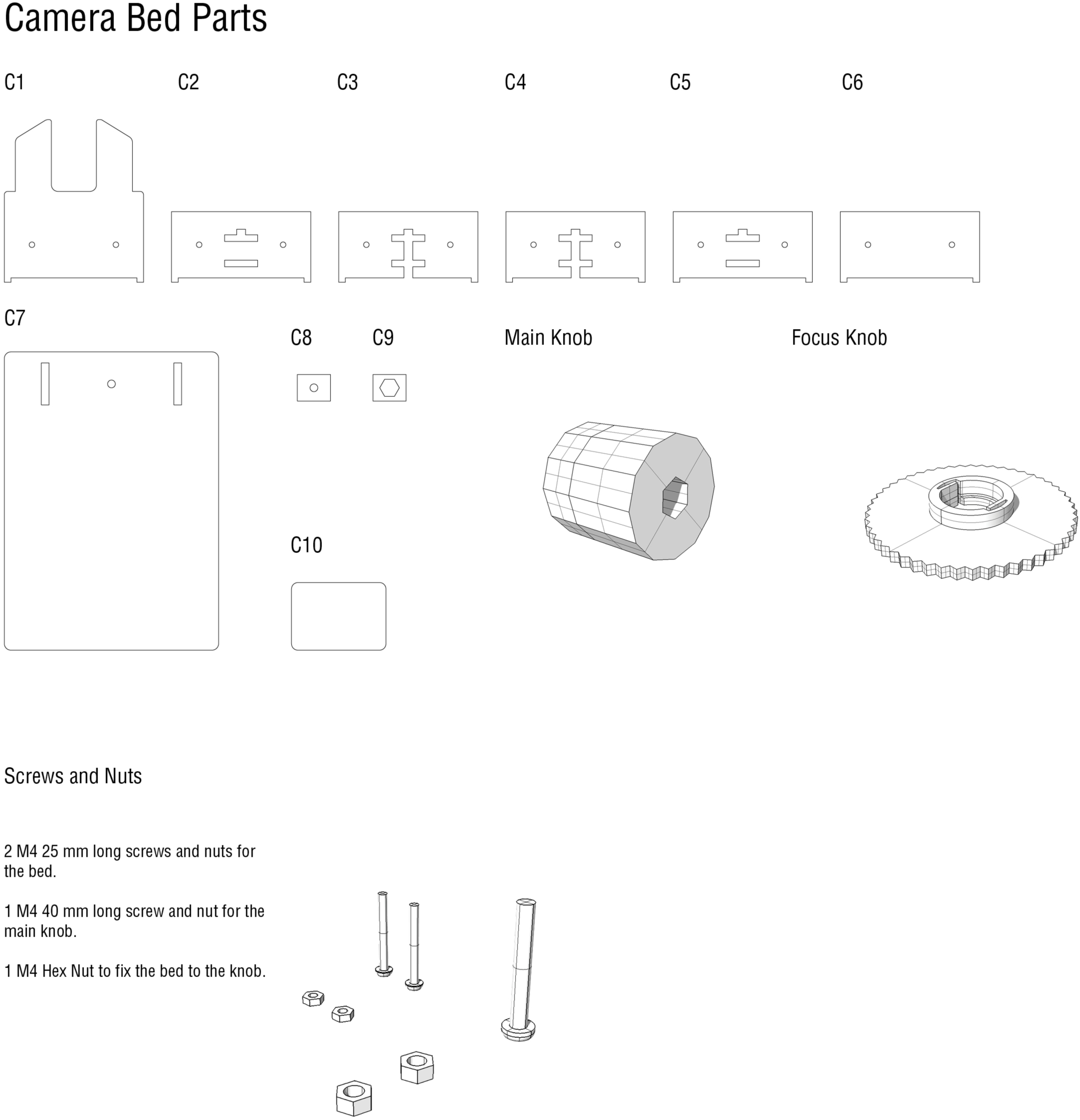

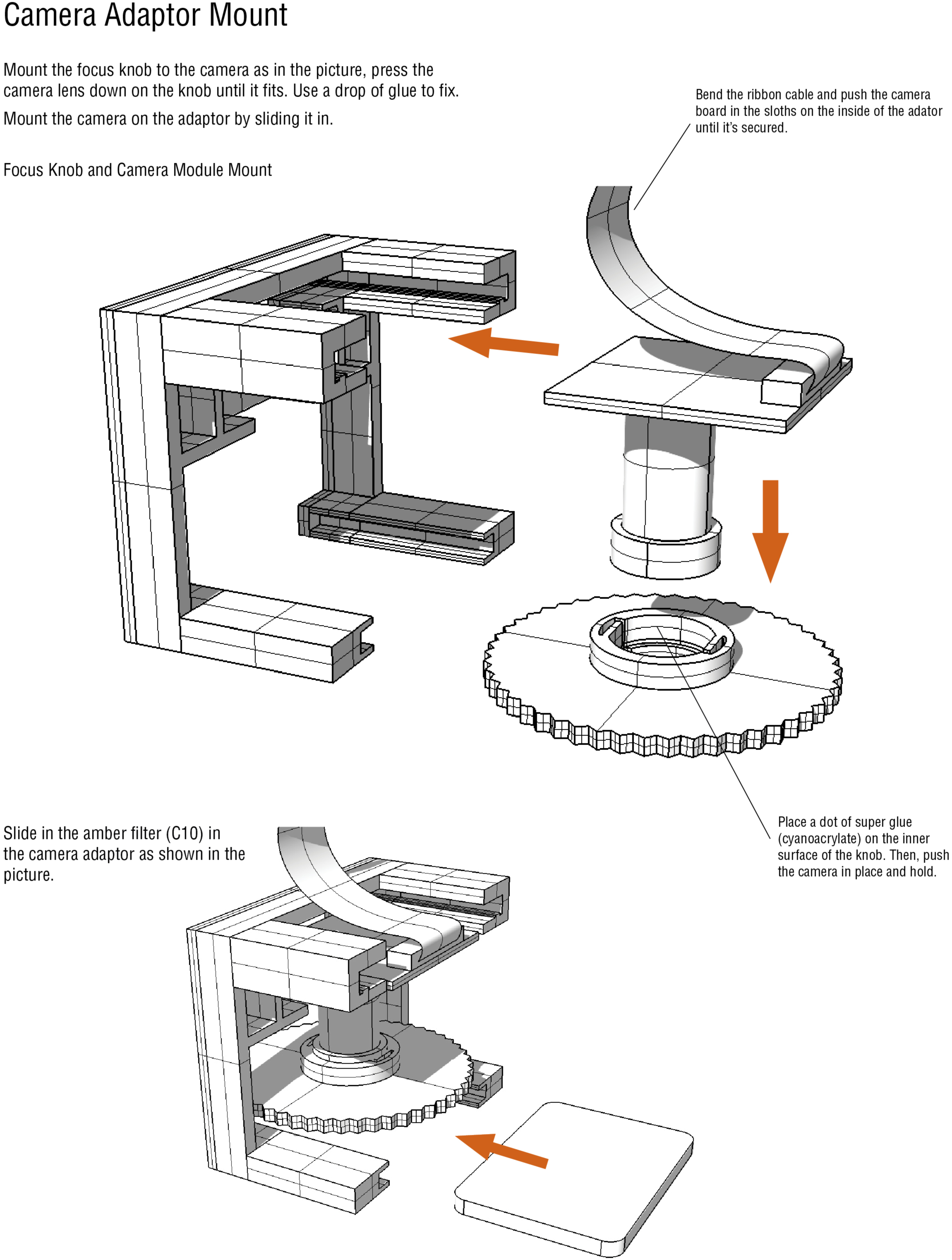

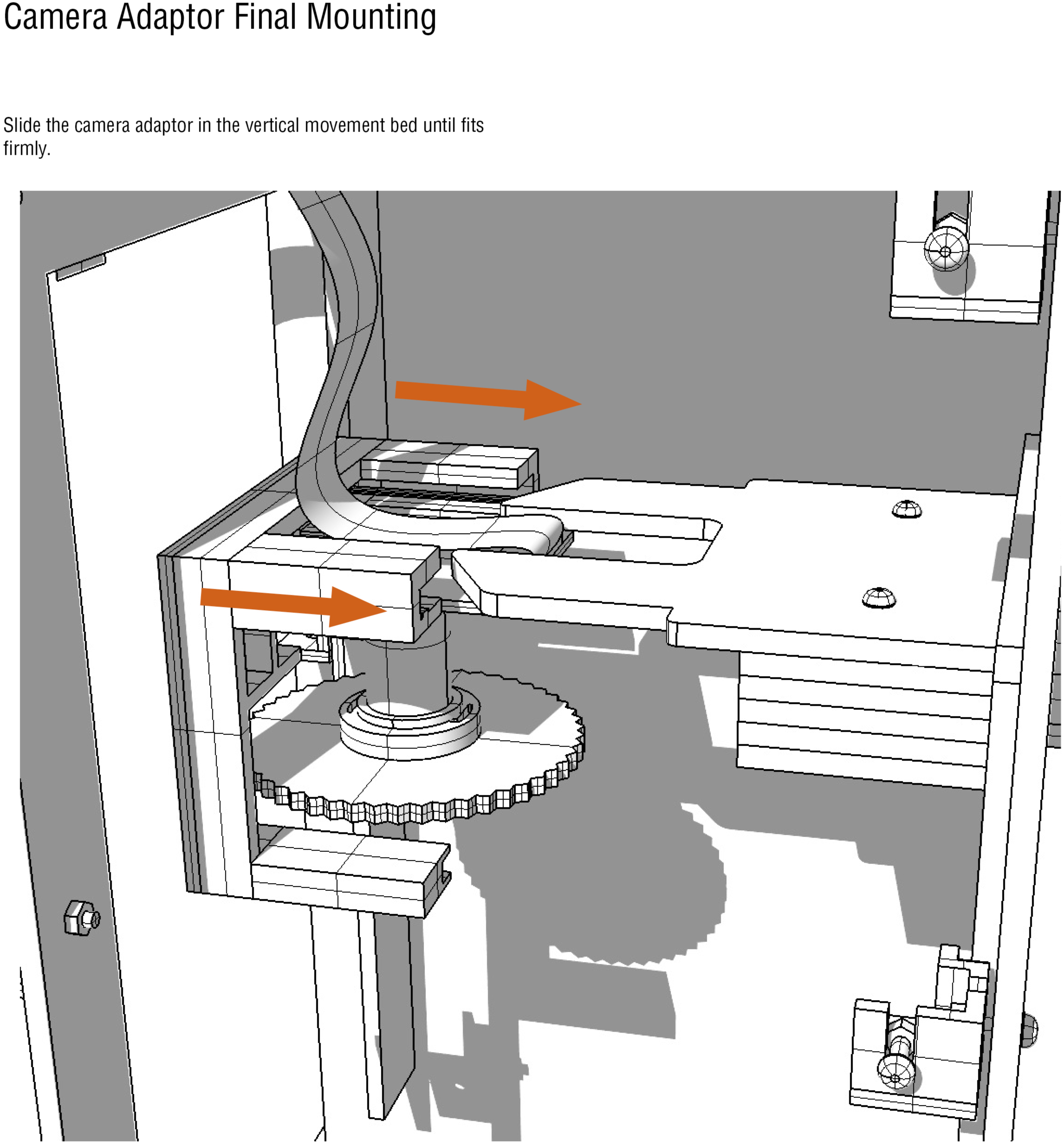

## S4 Appendix. assembly instructions for PCB

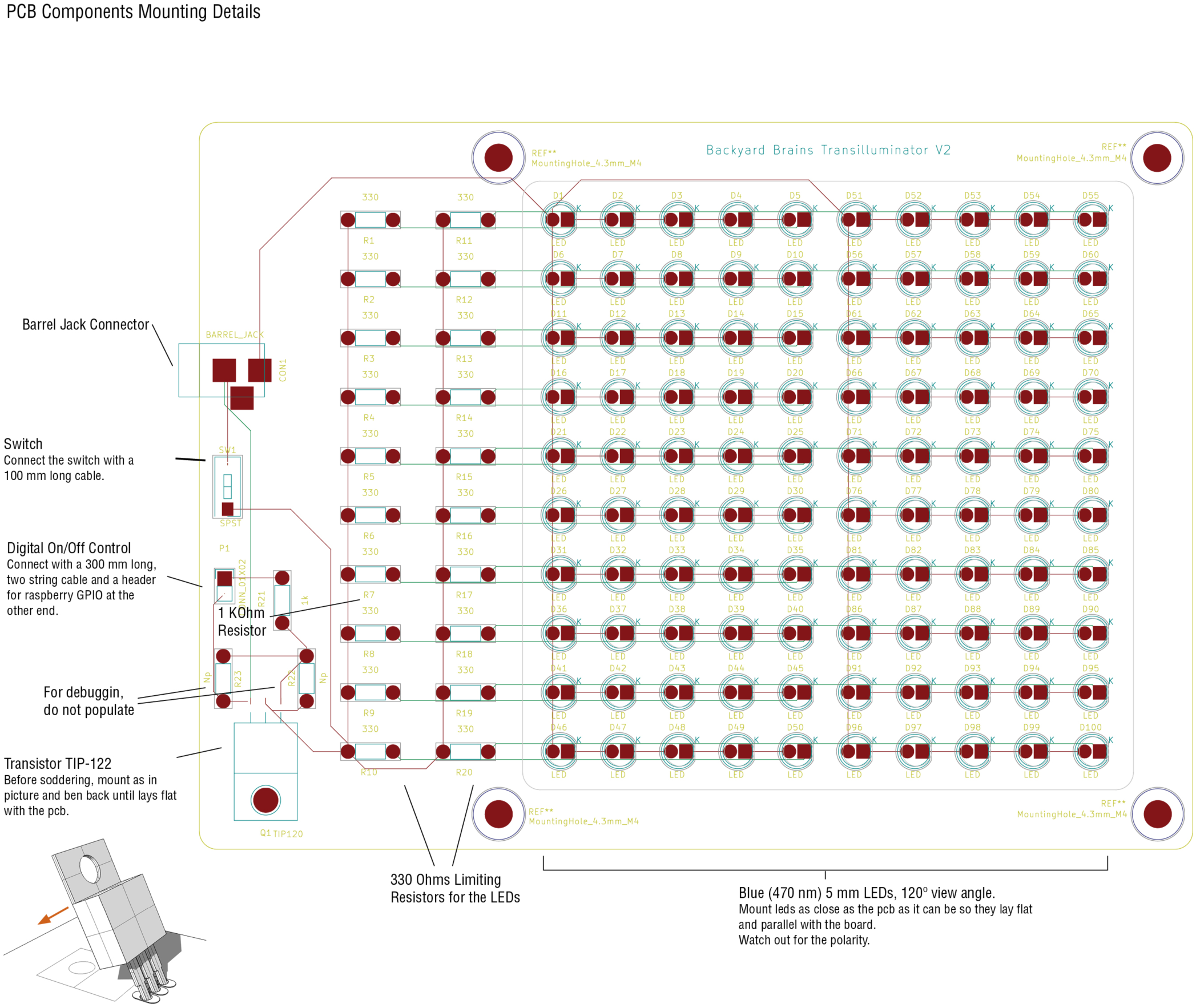

## S5 Appendix. Fluorescent signal mathematical analysis for time lapse images

### Fluorescence maths

August 16, 2017

#### 0.0.1 Fluorescent signal maths analysis for timelapse images

Intensity signal *I*_*s*_(*t*) can be decoupled on two terms: the signal given by the biological system *I*_*b*_(*t*) and the background signal given by the media *I*_*m*_(*t*).

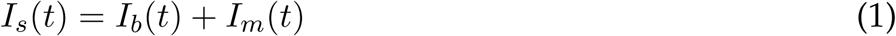

It is possible to obtain the background intensity directly from the data and get rid of him by simply substraction (assuming independence between them), focusing on the intensity given by the biological system.

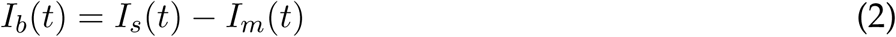

This signal intensity of interest (from here only “*I*(*t*)”) is given by the amount fluorescence retrieved by each cell. Then, it is possible to write the next expresión to represent this behavior:

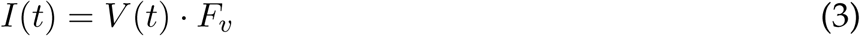

where *V* (*t*) is the volume of bacteria on each time point and pixel, and *F*_*v*_ is the pixel fluorescence emited by each unit of volume. As there are not precise measures of the colony volume or his absorbance at each point, would be useful to make some simplification. By assuming an homogeneus thickness along the colony (say h), then:

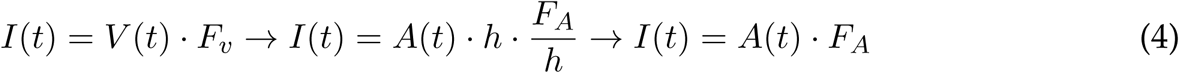

where *A*(*t*) is the area of bacteria on each time point and *F*_*A*_ is the amount of fluorescence emited per unit of area (pixel)

This last term could be assumed as a linear relation with the amount of fluorecent protein on each unit of area 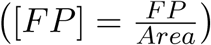:

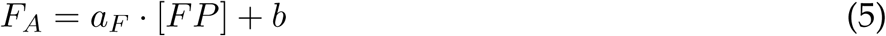

Then, the following expression is obtained for the intensity:

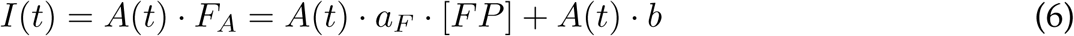

where the first term is given by the fluorescent protein amount on each cell and the second term is the bacterial colony autofluorescence.

Dividing by the area *A*(*t*) it’s is possible to obtain an expression for the mean fluorescent intensity per area:

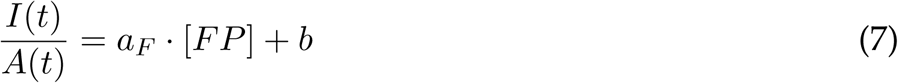

By taking the derivative over time, it’s get the expression:

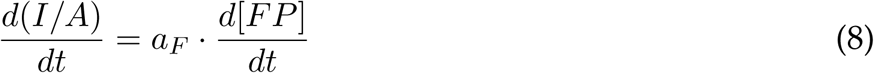

It is possible to relate the above with the dynamics of the colony protein expression. Taking simple models for transcription and traduction rate, respectively:

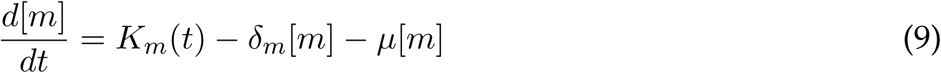

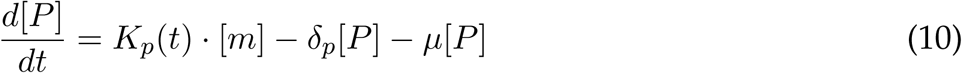

where *K*_*m*_(*t*) and *K*_*p*_(*t*) are the transcription and traduction rates, *δ* is the degradation rate and *μ* is the growth rate (being the last term the dilution one).

Under the assumptions of 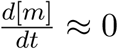 (and despising dilution) 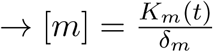, and taking *δ*_*p*_ *≈*0, the protein dynamics becomes:

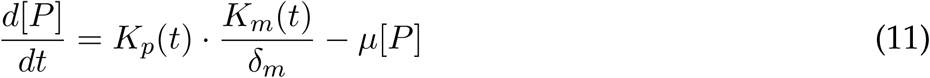

by grouping the protein expression term, it becomes:

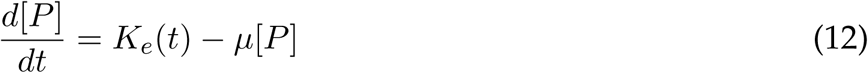

Using this model to represent the fluorescent protein dynamics, it’s possible use it on the mean fluorescence intensity expression developed (eq. 8):

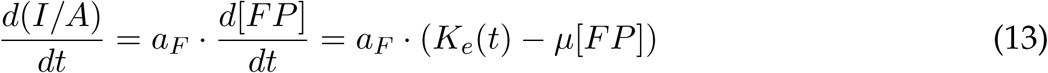

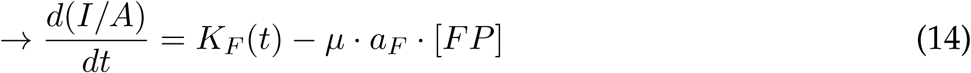

where *K*_*F*_ = *a*_*F*_·*K*_*e*_ is the flurescence intesity expression rate.

Finally, rearranging (14) and using (7):

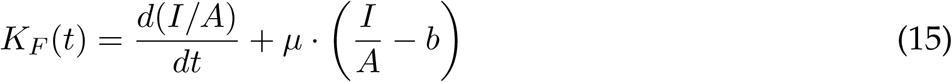

If despise the colony autofluorescence it’s possible to write:

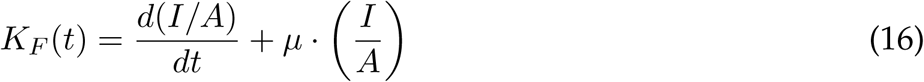

To write an expression for the mean intensity signal (*I/A*) and for the growth rate (*μ*), it’s possible to take the bacterial growth expression:

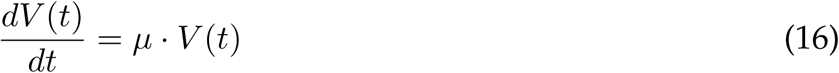

Which under assumption of constant thicknes it gets:

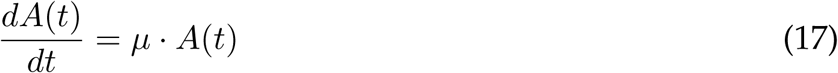

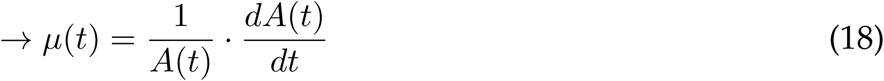

by taking a logistic area growth model:

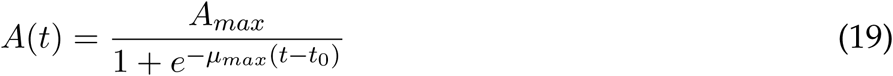

*μ*(*t*) gets:

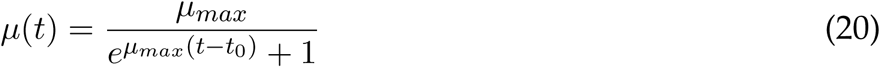

As we have colony area data, we can fit the model to them and compute *μ*(*t*) value. Also with the area values we can compute *I/A* directly from the data and smooth them with a spline (using scipy univariate spline) to compute the derivative of that function. Then we are able to compute the *K*_*F*_ (*t*) value on (15).

## S6 Appendix. Colony classification maths

Colony classificator maths

August 11, 2017

**0.1 Colony classificator maths**

Through approximating the relation between the fluorescent protein level with the fluorescence intensity as a linear one, it’s possible to write:
Fluorescence intensity vector = *A* · fluorescence protein level vector + basal signal

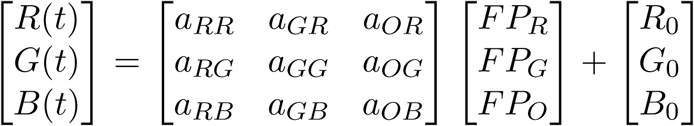

(where A is the linearization matrix that contains the slopes)

If it have only one fluorescent protein per colony, let’s say *F P_R_ >* 0, *F P_G_* = 0, *F P_O_* = 0, then:

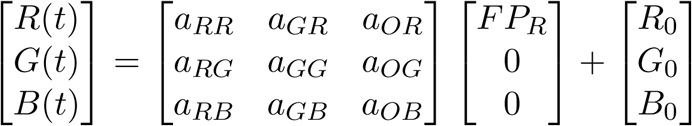

That is equal to:

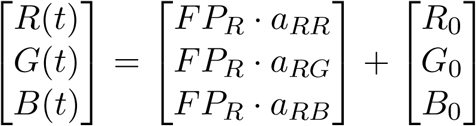

By taking only the red and green channels:

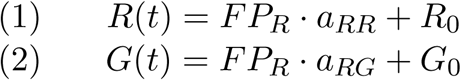

Assuming *a*_*RG*_ ≠0, rearrange (2) as:

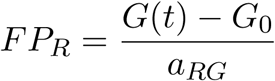

and replacing on (1):

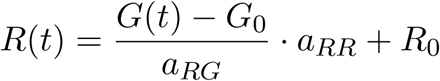

Rearranging it gives:

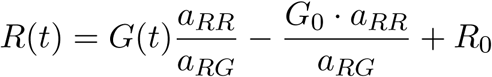

Finally, condensing the constants it becomes a linear relation between *R*(*t*) and *G*(*t*).:

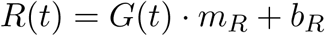

where 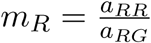 and 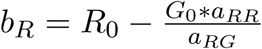

By proceeding the same way, it is possible to obtain a characteristic linear expression for each fluorescent protein:
when *FP*_*R*_ = 0, *F P_G_ >* 0, *F P_O_* = 0:

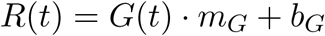

when *F P_R_* = 0, *F P_G_* = 0, *F P_O_ >* 0:

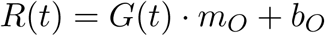

